# Higher temperatures worsen the effects of mutations on protein stability

**DOI:** 10.1101/2020.10.13.337972

**Authors:** Dimitrios - Georgios Kontopoulos, Ilias Patmanidis, Timothy G. Barraclough, Samraat Pawar

**Affiliations:** Science and Solutions for a Changing Planet DTP, Imperial College London, London, UK; Department of Life Sciences, Imperial College London, Silwood Park, Ascot, Berkshire, UK; Groningen Biomolecular Sciences and Biotechnology Institute, University of Groningen, Groningen, the Netherlands; Department of Zoology, University of Oxford, Oxford, Oxfordshire, UK

## Abstract

Understanding whether and how temperature increases alter the effects of mutations on protein stability is crucial for understanding the limits to thermal adaptation by organisms. Currently, it is generally assumed that the stability effects of mutations are independent of temperature. Yet, mutations should become increasingly destabilizing as temperature rises due to the increase in the energy of atoms. Here, by performing an extensive computational analysis on the essential enzyme adenylate kinase in prokaryotes, we show, for the first time, that mutations become more destabilizing with temperature both across and within species. Consistent with these findings, we find that substitution rates of prokaryotes decrease nonlinearly with temperature. Our results suggest that life on Earth likely originated in a moderately thermophilic and thermally fluctuating environment, and indicate that global warming should decrease the per-generation rate of molecular evolution of prokaryotes.

## Introduction

Genetic mutations are the primary source of the extensive phenotypic diversity that can be observed today, providing the raw material upon which selection can act. Thus, understanding the factors that influence the effects of mutations, as well as the rate at which they occur, is necessary for predicting the impacts of environmental change on organisms. From a thermodynamic standpoint, a given nonsynonymous mutation (i.e., a mutation that alters the amino acid sequence of a protein) can be classified as beneficial or detrimental based on its effect on the stability of the protein (***Zeldovich et al., 2007b***; ***Chen and Shakhnovich, 2009***; ***Quan et al., 2016***). Assuming a folding model with two states (unfolded and folded), we can define protein stability as the difference in Gibbs free energy (Δ*G* = *G*_unfolded_ − *G*_folded_). Nonsynonymous mutations alter Δ*G* by changing the free energy of the folded state. Therefore, the impact of a specific nonsynonymous mutation on protein stability can be calculated as ΔΔ*G* = Δ*G*_mutant_ − Δ*G*_wild_. Under this notation, mutations with positive ΔΔ*G* values are beneficial (stabilizing) and those with negative values are detrimental (destabilizing). Most mutations are destabilizing, with the average ΔΔ*G* close to −1 kcal/mol (***Zeldovich et al., 2007b***; ***Tokuriki et al., 2008***; ***Tokuriki and Tawfik, 2009***). Furthermore, an analysis of 2,188 point mutation experiments has shown that ΔΔ*G* values are almost independent of Δ*G*_wild_ (***Chen and Shakhnovich, 2009***), i.e., nonsynonymous mutations affect highly stable and less stable proteins quite similarly. For context, a mutation that leads to a Δ*G*_mutant_ value of 0 kcal/mol will result in half of the protein population being in the unfolded state (***Zeldovich et al., 2007b***; ***Chen and Shakhnovich, 2009***). This is expected to strongly impair the activity of the protein and can lead to the death of the organism, especially if the protein is essential for survival (***Zeldovich et al., 2007b***; ***Chen and Shakhnovich, 2009***).

The impact of temperature on Δ*G* is well-recognised and, for proteins that follow the two-state folding/unfolding model, the relationship between Δ*G* and temperature can be described by the Gibbs-Helmholtz equation (***Privalov 1990***; pink solid line in Figure 1; see also section I in Appendix 1). In contrast, there is no theoretical basis for the influence of temperature on the average ΔΔ*G*, possibly because it is extremely challenging to experimentally quantify the effects of all possible nonsynonymous mutations for a given protein at multiple temperatures. Temperature rises have often been reported to make mutations increasingly deleterious (temperature-sensitive mutations; ***Guo et al. 1996***; ***Dieterle et al. 2010***; ***Ndo et al. 2018***), but this is typically hypothesised to be exclusively driven by the decline in protein stability at high temperatures (***Puurtinen et al. 2016***; ***Berger et al. 2020***; Figure 1A). Under this hypothesis, ΔΔ*G* is assumed to be independent of temperature and constant (the “temperature-invariant ΔΔ*G*” hypothesis). Thus, mutations should become deleterious also at low temperatures where protein stability can be similarly low (***Puurtinen et al., 2016***).

**Figure 1.**
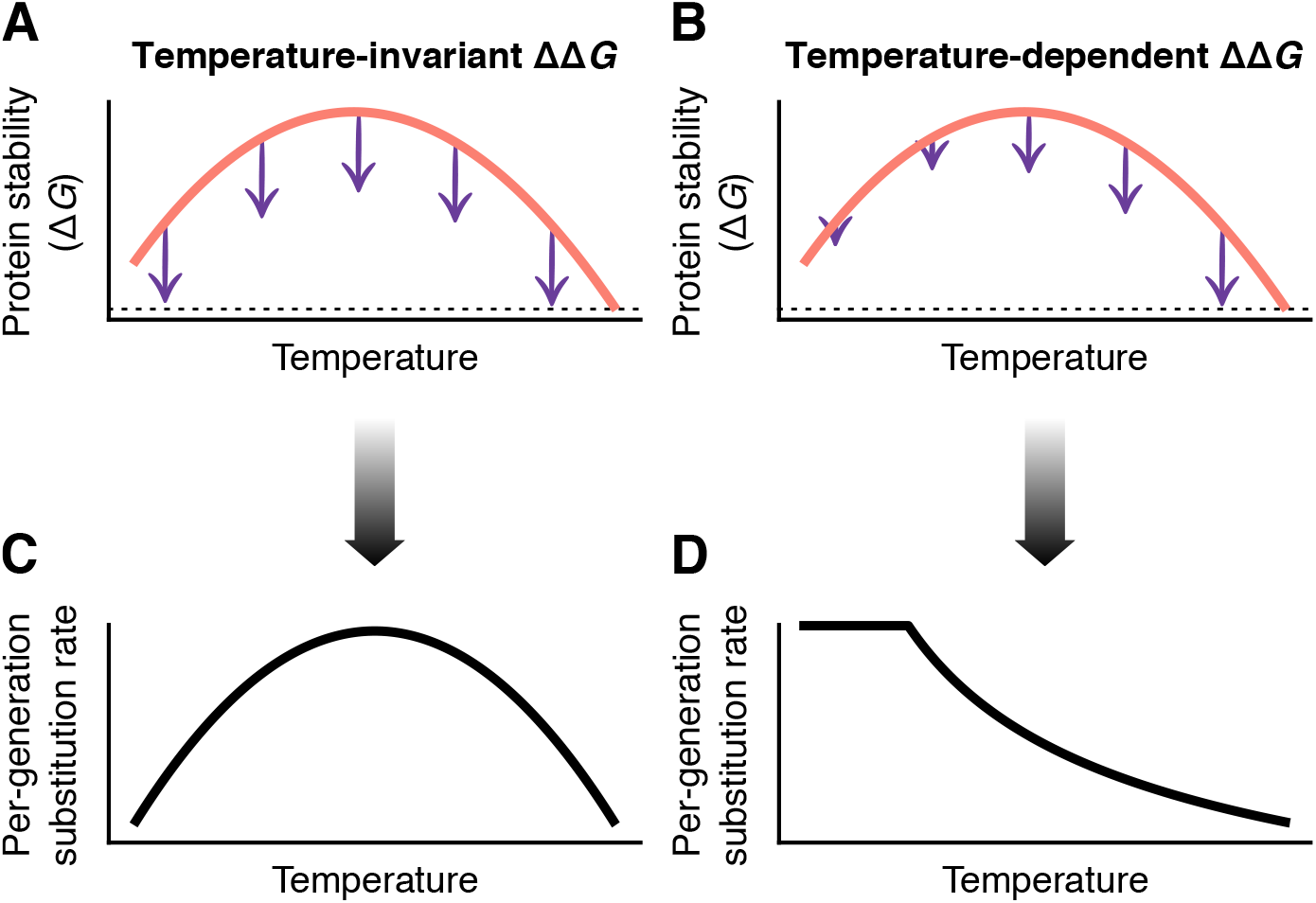
The two hypotheses tested in this study. The pink solid line represents the protein stability (Δ*G*) as a function of temperature, whereas the lengths of the purple arrows stand for the average effects of nonsynonymous mutations (ΔΔ*G*). Panel A represents the conventional assumption that mutational effects are independent of temperature, whereas in panel B, mutations become increasingly destabilizing as temperature increases. The dashed line indicates the position on the vertical axis of the critical threshold of Δ*G* = 0 kcal/mol, where the protein population is equally divided into the folded and unfolded states. The two hypotheses make different predictions for the relationship between per-generation substitution rate and temperature. In panel C, the substitution rate has a similar shape to that of the protein stability curve. In contrast, in panel D, the substitution rate is nearly invariant at low temperatures (where mutations have a negligible effect on protein stability), but decreases nonlinearly at higher temperatures.

In this study we introduce a new, alternative hypothesis (Figure 1B), that ΔΔ*G* is not constant but becomes more negative with temperature (the “temperature-dependent ΔΔ*G*” hypothesis). Such a relationship is expected from molecular biophysics. Consider a protein structure from a hyperther-mophilic organism which will have evolved under strong selection against heat denaturation, with this being reflected in its amino acid composition. In other words, the amino acid frequencies of a hyperthermophilic protein tend to differ from those of its psychrophilic or mesophilic orthologs (see e.g., ***Nakashima et al. 2003***; ***Zeldovich et al. 2007a***). At temperatures typically experienced by a hyperthermophile, a nonsynonymous mutation would (on average) strongly destabilize a key conformation of the protein, due to the high internal (kinetic and potential) energy of the atoms. In contrast, if the same mutation occurred at a lower temperature, atoms would have lower energy and the mutation would be better tolerated. Thermal shifts in the opposite direction (e.g., if a mesophilic protein is exposed to high temperatures) would have the opposite result (i.e., the mutation would become more detrimental). If this hypothesis holds, then there should be a negative interspecific relationship between environmental temperature and the average ΔΔ*G* value—estimated at a temperature close to those experienced by each species. A similar pattern would be expected between temperature and the average mutational effect within species.

Another difference between the two hypotheses lies in their predictions regarding the effects of temperature on the rate of molecular evolution (Figure 1C,D). Near the high temperature extreme, both hypotheses agree that mutations should bring the protein’s stability close to the critical threshold of 0 kcal/mol. Thus, thermophiles should have lower rates of sequence substitution, due to i) a higher fraction of deleterious mutations and potentially ii) the evolution of a low mutation rate through optimisation of DNA proofreading and repair mechanisms (***Kimura, 1987***). In cold environments, however, the temperature-invariant ΔΔ*G* hypothesis would also predict a low substitution rate (***Puurtinen et al., 2016***), as the resulting protein stability would be again close to the critical threshold (Figure 1A). In contrast, a low substitution rate in psychrophiles is not expected under the temperature-dependent ΔΔ*G* hypothesis (Figure 1B).

To test the two hypotheses, we used the essential enzyme adenylate kinase (ADK). This enzyme plays an important role in cellular energy homeostasis as it catalyses the biochemical reaction MgATP = AMP ⇌ MgADP = ADP, regulating adenine nucleotide levels in the cell. Studies in bacteria have shown that the stability of ADK is directly linked to fitness, suggesting the presence of strong selection (***Couñago and Shamoo, 2005***; ***Couñago et al., 2006***). We first collated a dataset of 70 ADKs of bacteria and archaea, spanning the entire temperature spectrum of life on Earth (from psychrophiles to hyperthermophiles; Appendix 1—table 1), and examined their evolutionary patterns. We then performed molecular dynamics simulations to obtain realistic conformations of each ADK at a temperature close to the natural environment of each species, and inferred the effects of all possible mutations (between 3,382 and 4,256 mutations per ADK depending on sequence length) on each conformation *in silico*. Lastly, we estimated the effects of temperature on i) the median ΔΔ*G* across and within species, and on ii) substitution rate (based on 44 genes).

## Results

### Dataset of ADK structures

We initially queried the Protein Data Bank (PDB) for experimentally determined ADK structures of bacteria and archaea. As this search yielded results from only 18 unique species (as of February 2018), we also performed homology modelling (***Brändén and Tooze, 1999***) to obtain a larger and more phylogenetically diverse dataset. Briefly, homology modelling allows for the inference of the 3D structure of a protein sequence (query sequence) based on similar protein sequences whose structures are experimentally determined (template sequences). To this end, we manually selected 70 bacterial and archaeal ADK sequences from the UniProt database (***The UniProt Consortium 2017***; Appendix 1—table 1), covering a broad range in terms of taxonomy (16 phyla; see Appendix 1—figure 4) and thermal preferences, and for which reliable homology models could be inferred using the Phyre2 web server (***Kelley et al. 2015***; see the Methods section). Regarding the thermal environment, in particular, we divided ADKs into four groups: psychrophiles (adapted to low temperatures and with an optimum temperature for growth ≤ 20°C), mesophiles (adapted to moderate temperatures and with an optimum temperature for growth between 20 and 40°C), thermophiles (adapted to high temperatures and with an optimum temperature for growth between 40 and 80°C), and hyperthermophiles (adapted to very high temperatures and with an optimum temperature for growth > 80°C). We tried to avoid ADKs from species which did not conform to these criteria (e.g., psychrotolerant species that are capable of growing at low temperatures but grow optimally at temperatures far above 20°C).

### 3D structure and evolution of prokaryotic ADKs

The ADK structure consists of three domains (Figure 2): the large and rigid CORE domain, the LID domain which binds ATP, and the NMPbind domain which binds AMP (***Li et al., 2015***). The length of the LID domain varies across species within our dataset, with most bacterial ADKs having a long LID domain and most archaeal ADKs a short one. Furthermore, archaeal ADKs with short LID domains have two additional *β*-strands in their CORE domain. These *β*-strands allow such ADKs to form homotrimers, whereas ADKs that lack them are monomeric (***Vonrhein et al., 1998***). The major conformational states that ADKs adopt are two: the open (inactive) conformation (Figure 2A,B) and the closed (active) conformation (Figure 2C,D). As the protein transitions between these two states, other intermediate conformations may be observed. In the absence of ligands, ADKs tend to shift between open and closed-like conformations, whereas ligand binding induces the closed conformation (***Kovermann et al., 2015***; ***Li et al., 2015***; ***Kovermann et al., 2017***).

**Figure 2.**
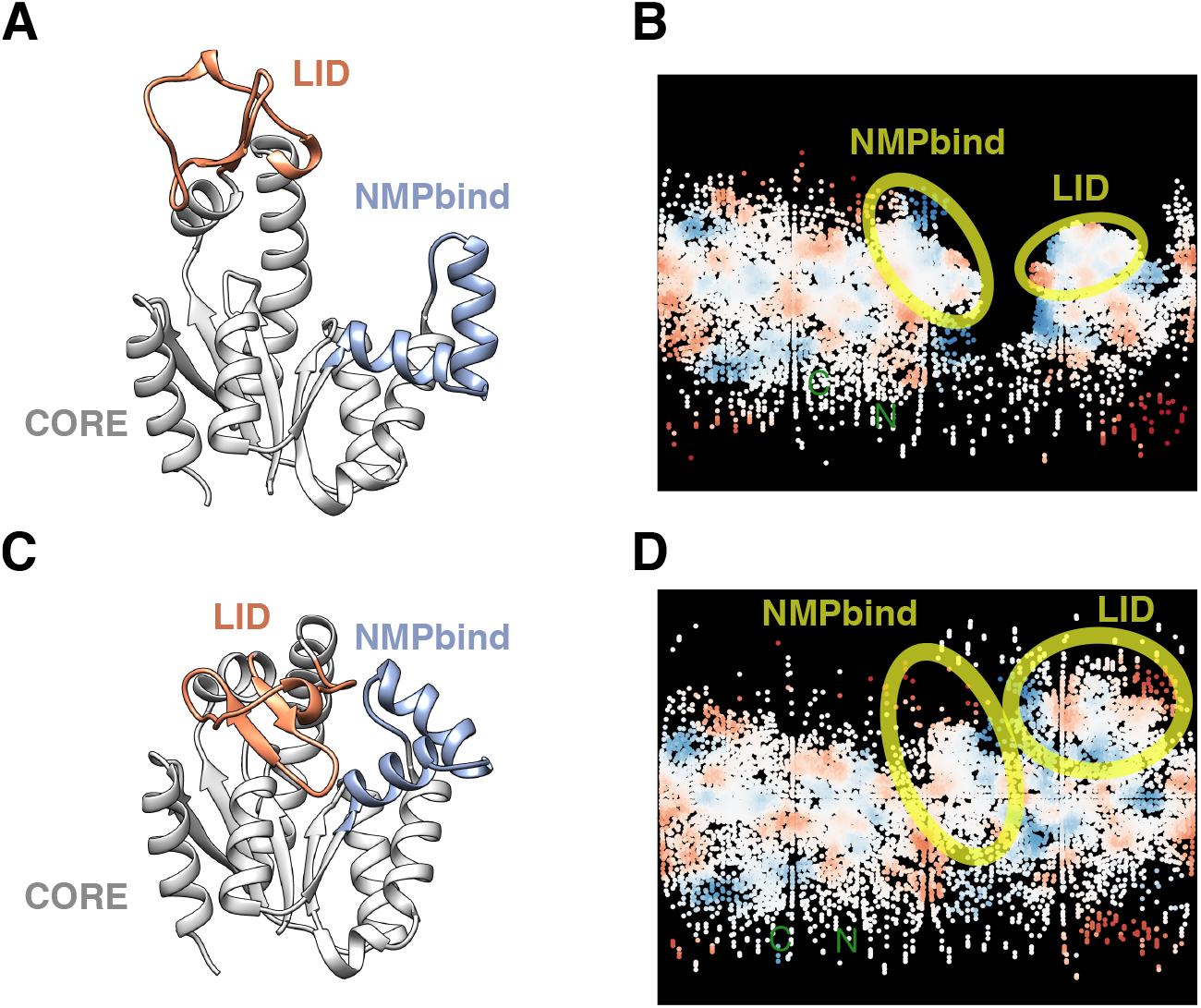
Open and closed conformations of the ADK of *Streptococcus pneumoniae* (PDB accession codes 4NTZ and 4NU0 respectively). The main structural domains are explicitly shown with different colors in panels A and C. Panels B and D are two-dimensional maps of the surface of each conformation, colored according to the charge of exposed amino acids (positively charged in blue, negatively charged in red). The open conformation (A,B) is energetically favourable in the absence of substrate. Upon substrate binding, the free energy of the closed conformation (C,D) becomes lower than that of the open conformation, and thus the former is preferred. A and C were rendered with UCSF Chimera v. 1.12 (***Pettersen et al., 2004***), whereas B and D with Structuprint v.1.001 (***Kontopoulos et al., 2016***).

The aforementioned structural differences among ADKs in our dataset (e.g., the length of the LID domain) may systematically affect the median ΔΔ*G* value, representing distinct evolutionary strategies against deleterious mutations. Thus, to better understand how variation in prokaryotic ADK structure emerges, we added an extra (outgroup) sequence (UniProt:Q9HKM7) to our dataset of 70 ADKs and reconstructed their evolutionary tree. The new sequence belongs to the archaeum *Thermoplasma acidophilum*, is described as a “putative adenylate kinase”, and is much shorter than any other ADK in our dataset. The 70 ADKs were found to form two major phylogenetic clusters (shown in purple and mustard colors; Figure 3A). The purple cluster contains structures from bacteria and archaea with both short and long LID domains, but without the two extra *β*-strands, causing these ADKs to form monomers (Figure 3B, subpanels I, II, and III). In contrast, the mustard cluster contains archaeal ADKs that can form homotrimers (Figure 3B, subpanel IV). Both clusters contain ADKs from two major archaeal clades: the Euryarchaeota and the TACK superphylum. This suggests that the two clusters are the outcome of a gene duplication that took place in the archaeal lineage, before the split of Euryarchaeota and TACK. The large distance that separates the two clusters, the divergence in secondary and quaternary structure between the two clusters, as well as systematic differences in their site-specific rates of evolution (inset of Figure 3A) are all consistent with this hypothesis. It is worth emphasising here that the activity of the ADK protein is directly linked to organismal fitness (***Couñago and Shamoo, 2005***; ***Couñago et al., 2006***). Thus, the significant structural differences that can be observed between the two ADK types would be highly unlikely to occur without the presence of two gene copies. One copy would remain functional, whereas the other would be able to evolve under relaxed selection pressure. Given that no species in our dataset expressed both ADK types (according to the UniProt database), either gene copy may have become a pseudogene or may have been removed through genome streamlining (***Ranea et al., 2005***; ***Giovannoni et al., 2014***). The hypothesis of a single gene duplication, followed by multiple gene losses is consistent with the phylogeny of the 70 species in our dataset, which we reconstructed from 44 genes (Figure 3C; see Methods and section IV in Appendix 1). Nevertheless, it is also possible that the distribution of the two ADK types across species may be partly the outcome of horizontal gene transfer events. In any case, we henceforth use the terms “ancestral-type” to refer to ADKs from the purple cluster and “archaeal-type” to refer to those from the mustard cluster.

**Figure 3.**
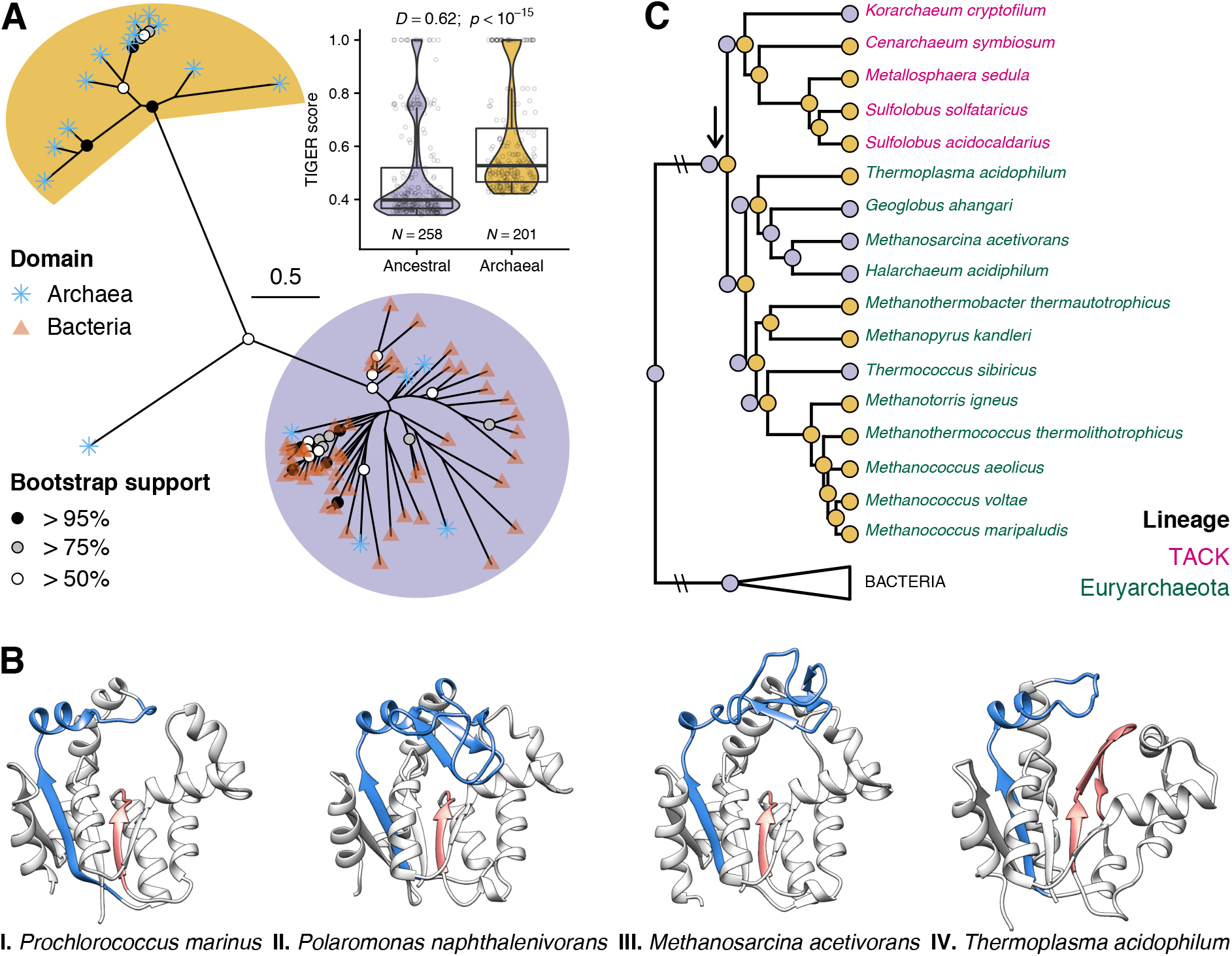
Evolution of ADKs of bacteria and archaea. A: Prokaryotic ADKs are broadly divided into two phylogenetic clusters, possibly as a result of an early gene duplication event. The ADKs of the purple (ancestral-type) cluster are monomeric, whereas those of the mustard (archaeal-type) cluster are homotrimeric. The archaeal sequence that belongs to neither cluster is the outgroup sequence. Inset: Distributions of TIGER scores (a proxy for site-specific evolutionary rate; ***Cummins and McInerney 2011***) for the ADKs of the two phylogenetic clusters, which we aligned separately. Boxplot edges represent the first and third quartiles, whereas the solid line stands for the median. Whiskers extend up to the most remote data point within 1.5 interquartile ranges from each boxplot edge. The difference between the two TIGER score distributions is statistically significant according to the two-sided two-sample Kolmogorov-Smirnov test (***Corder and Foreman, 2014***). Their low median values highlight that both ADK types (but predominantly those that are monomeric) exhibit very high rates of evolution. This explains the poor statistical support (lack of black or grey circles) of most tree nodes. B: Four representative ADK structures from bacteria (subpanels I and II) and archaea (subpanels III and IV). The main differences among them (i.e., LID length and presence/lack of two additional *β*-strands) are highlighted in blue and red. Structures I-III are part of the purple phylogenetic cluster, whereas IV is part of the mustard cluster. C: The proposed hypothesis regarding the emergence of the two ADK types (shown as purple and mustard circles). The arrow indicates the position of the hypothesised gene duplication event. Ancestral species (tree nodes) that presumably had both ADK types are shown with both circles.

### Mutations become more destabilizing with temperature across and within species

Next, we examined if the median effect of all possible mutations on each ADK varies with environmental temperature across species. To this end, we conducted molecular dynamics simulations (200 ns long, with 10 independent replicates) for each ADK at a temperature close to those experienced by each species: 6.85°C for psychrophiles, 26.85°C for mesophiles, 56.85°C for thermophiles, and 81.85°C for hyperthermophiles. These simulations enabled us to a) obtain reliable conformations for each ADK at its biologically meaningful temperature, and to b) remove any minor structural biases arising from homology modelling using multiple templates. Sampled ADK conformations were grouped into clusters (from 2 to 20), and an average (representative) structure was calculated for each conformational cluster (see Methods and Appendix 1—figure 7 for an example). We then submitted each representative structure to the STRUM web service (***Quan et al., 2016***), a machine learning tool that has been trained and tested on experimentally-determined ΔΔ*G* values from the ProTherm database (***Kumar et al., 2006***). STRUM can provide reasonably accurate ΔΔ*G* estimates for all possible single point mutations of a given protein, based on sequence and structure information. After obtaining estimates of the median ΔΔ*G* value per ADK conformation, we calculated the weighted median mutational effect across all conformations 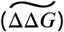, with weights equal to the proportion of each conformational cluster across simulations.

We checked the 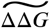 values for two possible biases: the type of the conformation (open versus closed) and the quality of the initial homology model. For the former, we used the R package bio3d (v. 2.3-4; ***Grant et al. 2006***) to calculate the radius of gyration of each conformation, which is a measure of protein structure compactness. For the latter, we used the proportion of amino acids in disallowed regions (see the Methods section) as a metric. No systematic association could be detected between 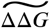 and these two factors (Appendix 1—figure 8).

To investigate if 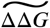 varies with temperature, we fitted a number of alternative regression models with the MCMCglmm R package (v. 2.26; ***Hadfield 2010***; see the Methods section). We found that 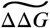 becomes linearly more negative (mutations more destabilizing) with temperature (Figure 4A, Appendix 1—table 4). The two ADK types share a common slope for temperature but have different intercepts, with archaeal-type ADKs being more robust to nonsynonymous mutations than ancestral-type ADKs. A systematic effect of LID length on 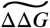 could not be detected, and controlling for phylogeny did not improve the fit either (Appendix 1—table 4).

**Figure 4.**
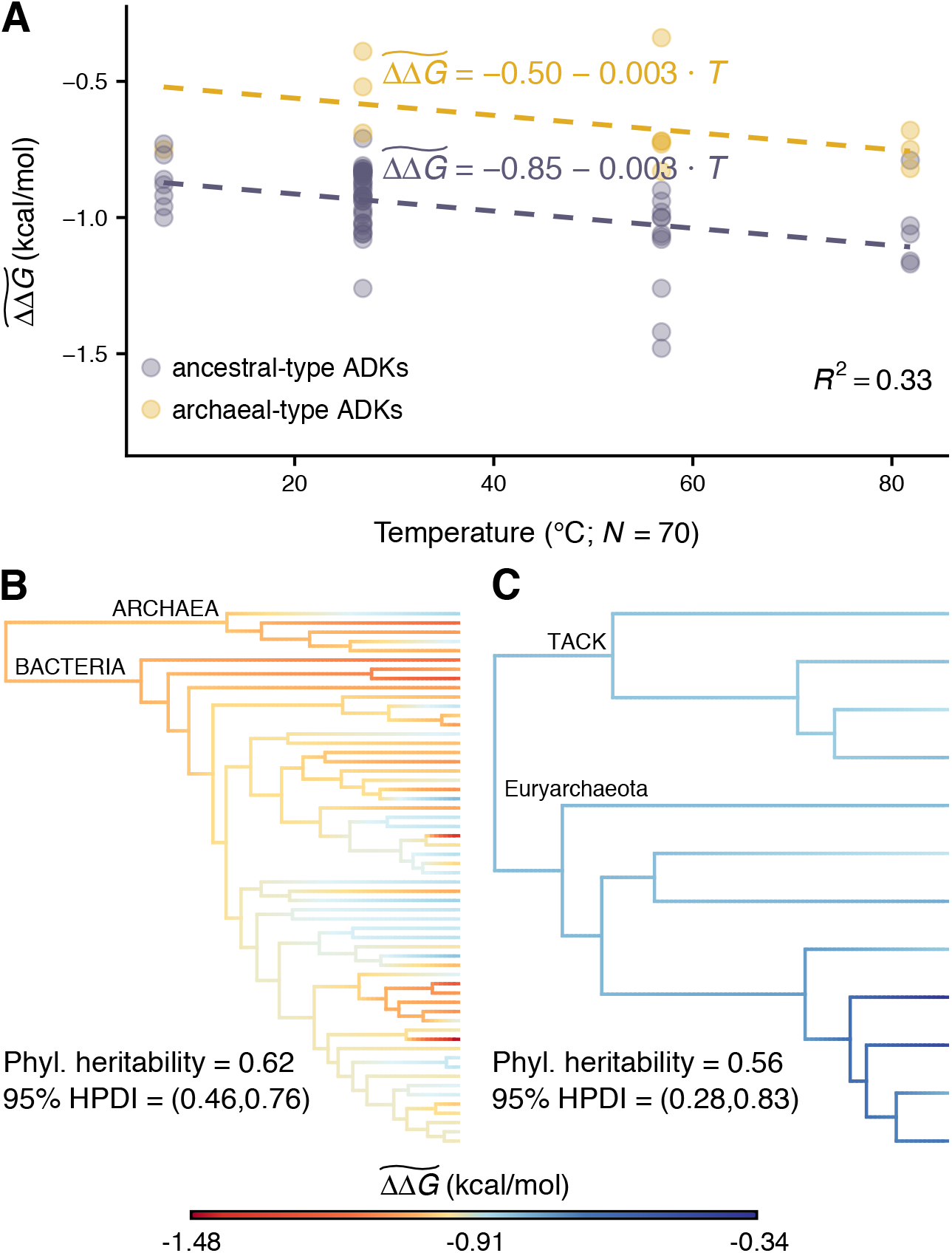
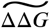 varies systematically with temperature and is phylogenetically heritable. A: Across species, nonsynonymous mutations become on average more destabilizing with temperature. Each data point represents the weighted median effect of all possible mutations for a single ADK at a temperature close to those experienced by the species. Mutations are less detrimental to archaeal-type ADKs, suggesting a potential evolutionary benefit for species whose genomes code for this ADK type. All coefficients of the model had 95% Highest Posterior Density (HPD) intervals that did not include zero. B-C: For both ancestral (panel B) and archaeal (panel C) ADK types, 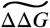 evolves along the species phylogeny, reflecting the evolution of species’ thermal strategies (e.g., adaptation to psychrophilic temperatures). Nevertheless, the phylogenetic heritability is lower than that expected by a random walk in the parameter space (Brownian motion), possibly in part due to episodes of convergent evolution or interspecific recombination of the ADK gene (***Feil et al., 1996***). 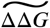 values of internal nodes were estimated using the stable model (***Elliot and Mooers, 2014***), which can accommodate patterns of both gradual and rapid trait evolution. Panels B and C were rendered using the phytools R package v. 0.6-60 (***Revell, 2012***).

Further support for a non-constant 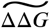 was obtained through the estimation of its phylogenetic heritability, i.e., the extent to which closely related species (which often tend to live in similar thermal environments) have more similar 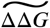 values than randomly selected species. A complete absence of phylogenetic heritability would be expected if the average effect of mutations was effectively constant across temperatures and species. In contrast, for both ADK types, 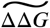 exhibited intermediate phylogenetic heritability. This indicates that it evolves along the phylogeny but not in a purely random and gradual (Brownian motion) manner (Figure 4B-C), likely reflecting the rapid evolution of the ADK sequence (see the inset of Figure 3A).

We additionally checked if the temperature-dependent ΔΔ*G* hypothesis also holds within species. For this, we selected five ancestral-type ADKs (for consistency reasons) from our dataset: i) from *Aquifex aeolicus* (a hyperthermophilic bacterium), ii) from *Thermus thermophilus* (a thermophilic bacterium), iii) from *Mycobacterium tuberculosis* (a mesophilic bacterium), iv) from *Sporosarcina globispora* (a psychrophilic bacterium), and v) from *Methanosarcina acetivorans* (a mesophilic archaeum). We also performed ancestral sequence reconstruction and homology modelling to infer the ADK of the last universal common ancestor (LUCA; section VIII in Appendix 1). An analysis of the resulting sequence showed that the enzyme’s optimum temperature for catalysis (*T*_opt_) is estimated at 65.23°C (95% prediction interval = [54.79, 75.67]). Given that *T*_opt_ values of prokaryotic ADKs tend to fall close to the temperature of maximum growth rate (Appendix 1—figure 12), this result suggests that the LUCA likely experienced weakly thermophilic conditions.

We then performed molecular dynamics simulations for each of these 6 ADKs at 8 different temperatures, from 6.85°C to 94.35°C. This allowed us to infer the effects of mutations on each ADK across a wide temperature range, including temperatures in which the species would not be able to survive. As previously, representative structures were submitted to STRUM to obtain estimates of their median ΔΔ*G*. Finally, we used weighted least squares regression to fit six mathematical equations (Appendix 1—figure 9) to the relationship between median ΔΔ*G* and temperature of each species. The best-fitting equation was determined according to the small sample size-corrected Akaike Information Criterion (***Sugiura 1978***; AICc). For all ADKs, median ΔΔ*G* becomes more negative with temperature (Figure 5), as expected. It is also worth noting that the shape of the relationship between median ΔΔ*G* and temperature varies to an extent across ADKs of different species (see also Appendix 1—figure 10).

**Figure 5.**
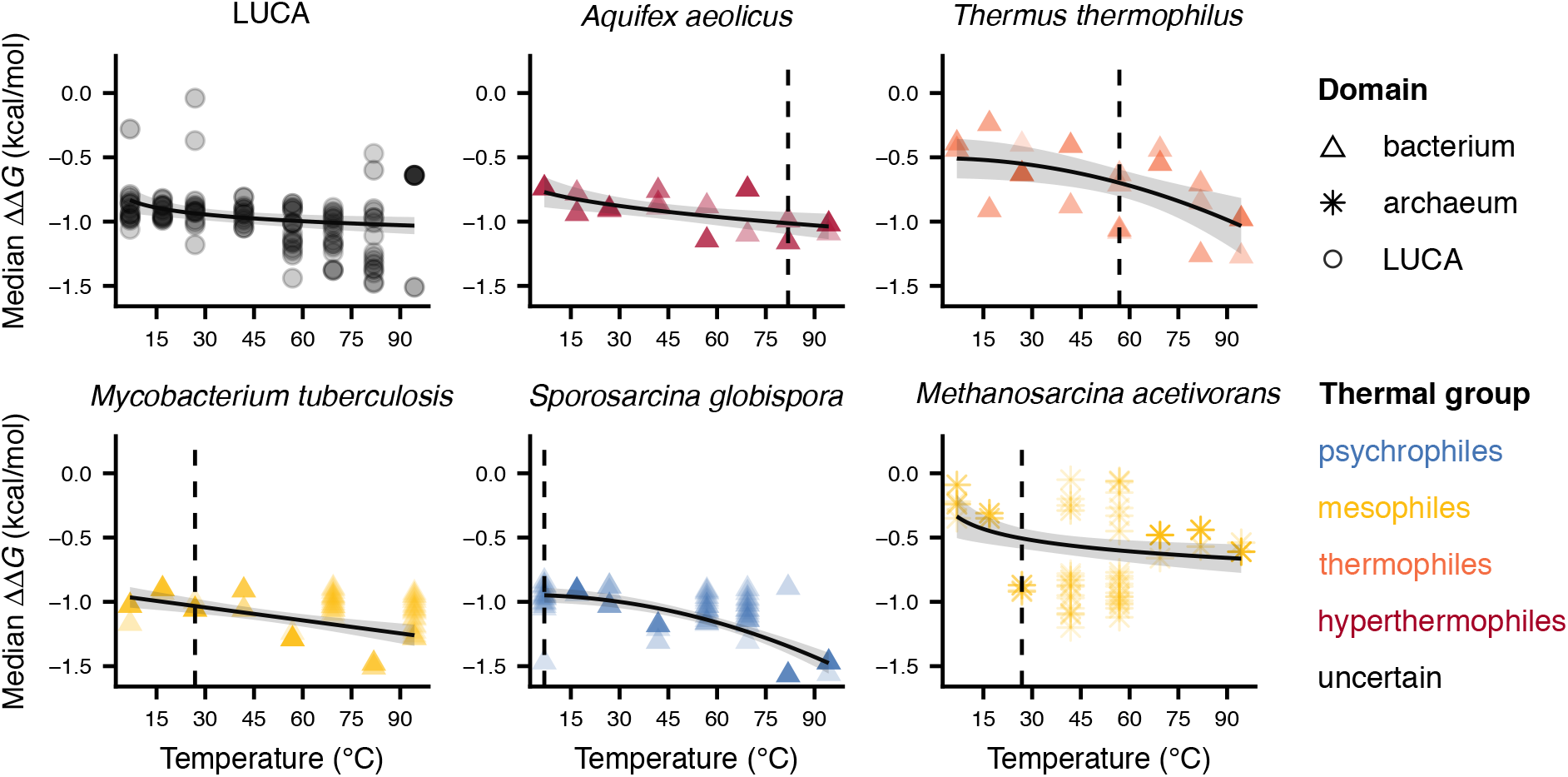
The relationship of median mutational effects versus temperature for the ancestral-type ADKs of five extant species and of the Last Universal Common Ancestor. Each data point corresponds to a different representative conformation, whereas the color depth stands for the proportion of protein snapshots per conformation at a given temperature. The vertical line shows a temperature near those typically experienced by each species (i.e., the temperatures shown in Figure 4A). The observed relationships vary both in their position along the vertical axis and in their sensitivity to increasing temperature.

### Substitution rates of prokaryotes decrease with temperature

Finally, we tested the predictions of the two hypotheses of Figure 1 for the relationship between substitution rate and temperature across the 70 species in our ADK dataset. It is worth noting that this relationship may be confounded by other (potentially temperature-dependent) factors such as cell volume or generation time (see sections X and XI in Appendix 1). Therefore, to control for any effects of these two factors on substitution rate, we used the following equation (see section XI in Appendix 1):

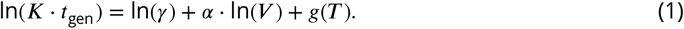

Here, *K* is the substitution rate per time, *t*_gen_ the generation time, *T* temperature, *V* the cell volume, *γ* a normalization constant (independent of cell volume and temperature), *α* a dimensionless exponent, whereas *g*(*T*) is a temperature dependence function. To accommodate the predictions of both hypotheses of this study (Figure 1), we specified *g*(*T*) as either a logarithmic or a quadratic equation. Given that we had previously reconstructed and calibrated a 44-gene phylogeny of the species in our dataset (see Methods and section IV in Appendix 1), we had already obtained estimates of the substitution rate for each branch of the phylogeny. Thus, we used the median *K* estimate of the terminal branch corresponding to each of the 70 species. For temperature (*T*), we assigned to each species the temperature value for its group (i.e., 6.85°C for psychrophiles, 26.85°C for mesophiles, 56.85°C for thermophiles, and 81.85°C for hyperthermophiles). Regarding generation time (*t*_gen_) estimates, we analysed a large dataset of prokaryotic growth rates (***Smith et al., 2019***) and obtained a general equation that links generation time with environmental temperature (section X in Appendix 1). Lastly, cell volume estimates were calculated from linear cell dimension measurements which we collected from the literature for all but one species (section IX in Appendix 1).

We then fitted a series of alternative regression models with MCMCglmm and performed model selection to identify the most appropriate model (Appendix 1—table 6). Through this process, we detected a strong negative association between generation time-corrected substitution rates and temperature (Figure 6). In contrast, a systematic effect of cell volume on substitution rate was not found, as the exponent of cell volume was statistically indistinguishable from zero.

**Figure 6.**
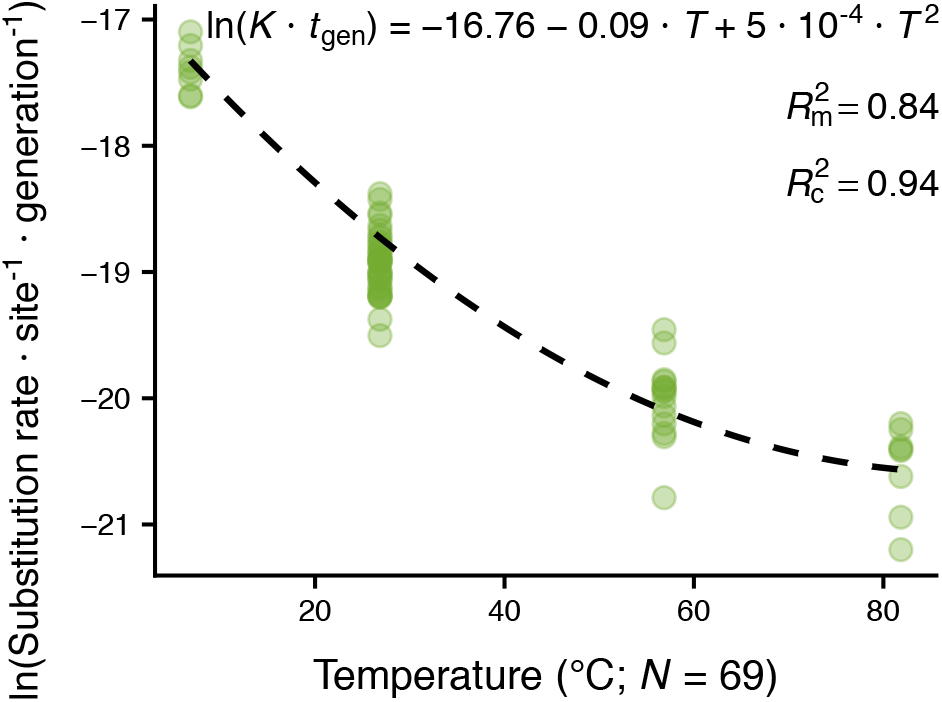
The generation time-corrected substitution rate decreases with temperature. This nonlinear monotonic relationship is consistent with our finding that mutational effects are temperature-dependent, becoming increasingly deleterious as temperature increases. Temperature explains 84% of variance on its own, whereas an additional 10% of variance is explained by a phylogenetic random effect on the intercept.

## Discussion

Nonsynonymous mutations have hitherto been considered to become more deleterious with temperature exclusively because of the decline in protein stability (Δ*G*) beyond a critical temperature (Figure 1A). Our study shows that besides temperature-driven changes in protein stability, mutations themselves (ΔΔ*G*) become on average increasingly destabilizing as temperature increases (Figure 1B). This aggravation of the effects of mutations on protein stability is consistent with the increased energy of atoms at high temperatures.

In particular, our interspecific analysis of the weighted median effects of all possible mutations 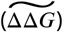 across two types of prokaryotic ADKs revealed that 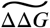 decreases linearly with environmental temperature and evolves across species (Figure 4), despite the relatively limited phylogenetic signal present in the alignment of ADK sequences (Figure 3A). Furthermore, ADK-specific relation-ships between median ΔΔ*G* and temperature were negative (as expected) and varied predominantly in their elevation along the median ΔΔ*G* axis (vertical axis of Figure 5 subpanels, Appendix 1—figure 10), and to a lesser extent in their temperature sensitivity. Variation in temperature sensitivity is likely to reflect the conditions of the local environment, as thermal generalist species or strains (with low temperature sensitivity) should be selected for at environments with large intergenerational temperature fluctuations (***Gilchrist, 1995***). In particular, given that the LUCA had a relatively low temperature sensitivity, we would expect it to have emerged in a strongly fluctuating thermal environment. Indeed, currently proposed environments for the origin of life (e.g., alkaline hydrothermal vents, subaerial hot springs, and submarine hydrothermal sediments; ***Mulkidjanian et al. 2012***; ***Sojo et al. 2016***; ***Westall et al. 2018***; ***Camprubí et al. 2019***) are characterised by strong gradients in temperature, pH, and other physicochemical variables. Such environments are also consistent (under certain assumptions) with the LUCA being adapted to a moderate thermophilic (but not hyperthermophilic) lifestyle, as suggested by the relatively high robustness to mutations of its ADK, its estimated *T*_opt_ for catalysis (65.23°C; Appendix 1—figure 12), as well as some previous studies (***Di Giulio, 2003***; ***Gaucher et al., 2003***, ***2008***; ***Catchpole and Forterre, 2019***).

Consistent with all the above results, we detected a monotonic nonlinear decrease of the per-generation substitution rate (averaged across 44 genes) with temperature (Figure 6). According to the neutral theory of molecular evolution (***Kimura, 1987***), such a pattern could occur if i) the proportion of deleterious mutations increases with temperature (as we show in this study) and/or if ii) mutation rates decrease with temperature. An investigation of the latter cannot be done without new laboratory experiments, given that almost all available prokaryotic mutation rate measurements are from mesophiles (Appendix 1—figure 16). A third factor that may partly drive the decline in substitution rates with temperature is the effective population size. More precisely, in species with low effective population size, weakly deleterious mutations may not be removed by selection, yielding an elevated substitution rate (***Woolfit, 2009***). Thus, if the effective population size increases with temperature across prokaryotes, this would partly explain the high substitution rates of psychrophiles. Once again, currently existing datasets are inadequate for addressing this hypothesis.

To the best of our knowledge, such systematic inter- and intraspecific relationships between the median mutational effect and temperature have never been reported before in the literature. This is likely explained by the complete lack of experimentally-determined datasets of ΔΔ*G* (averaged across all possible mutations) for a given protein at multiple temperatures. The generality of these relationships could be investigated by performing similar analyses for other kinases (or other proteins in general) in future studies. If, for example, the slope of the interspecific relationship varies very little across different proteins, it could suggest the presence of a hard biophysical constraint on thermal adaptation. Similarly, determining the temperature sensitivity of the median ΔΔ*G* value for many ADKs across thermal gradients could shed more light on the likely effects of the local environment.

Our finding of a decline in substitution rates with temperature appears to contradict the results of two previous studies by Gillooly and co-workers (***Gillooly et al., 2005***, ***2007***). These studies found that across animals, nucleotide and amino acid substitution rates increase with temperature and decrease with body mass, consistent with a dependence of mutation rate on metabolic rate. The discrepancy between their results and ours probably arises from two main differences in study design. First, Gillooly and co-workers assumed that generation time has the same temperature and body size dependence as mass-specific metabolic rate. Second, because they only examined the substitution rates of animals, their datasets did not cover the entire temperature range, but mainly mesophilic temperatures.

Overall, the present study sheds new light on the mechanism by which nonsynonymous mutations become increasingly deleterious as temperature increases. Our results are consistent with and help explain the monotonic decrease in the per-generation substitution rates of prokaryotes with temperature, across a 75-degree temperature range. These findings further our understanding of the mechanisms underlying thermal adaptation in prokaryotes, from the origin of life to the present day, improving—among others—our ability to predict the potential of populations and species to develop evolutionary responses to ongoing global change.

## Methods

### Homology modelling

We performed homology modelling using the intensive mode of the Phyre2 server which can produce a homology model based on multiple templates available from the PDB. The choice of the most appropriate template(s) is made by taking into account i) the confidence of the alignment between the query and template sequences, and ii) the coverage of the query sequence by the template(s). Any amino acids of the query sequence that are not covered by the template(s) are modelled using an *ab initio* approach. As the accuracy of *ab initio* modelling is usually lower, we rejected any homology models for which more than four amino acids had to be modelled *ab initio*.

We also ensured that the sequence identity between each query sequence and its chosen template(s) was sufficiently high. For this, we used Rost’s empirical equation (***Rost, 1999***):

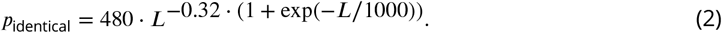

*p*_identical_ is the percentage of sequence identity and *L* the length of the alignment. Plotting this equation for different values of *L* gives a line that partitions the parameter space into a “safe zone” for homology modelling (i.e., where the resulting structural models are expected to be reliable) and a “twilight zone” (Appendix 1—figure 2).

Finally, we performed a Ramachandran plot (***Brändén and Tooze, 1999***) analysis to assess the quality of the resulting homology models. The two axes of the Ramachandran plot correspond to the backbone dihedral angles *ϕ* and *ψ* of amino acids of a given protein structure. Based on a large number of experimentally determined structures, the parameter space defined by the two axes has been empirically partitioned into the “most favourable regions”, “additional allowed regions”, “generously allowed regions”, and “disallowed regions”. Thus, the amino acids of a high quality model are expected to be mostly found in the most favoured and additional allowed regions, with none or very few amino acids in disallowed regions. We generated Ramachandran plots for each ADK structure with PROCHECK (***Laskowski et al., 1993***) and examined the proportions of amino acids in each region of the plot (Appendix 1—figure 3).

### Phylogenetic analyses

#### ADK tree inference

ADK sequences were aligned with MAFFT (v7.123b; ***Katoh and Standley 2013***) using the L-INS-i algorithm. We then ran ModelTest-NG (***Darriba et al., 2020***) to identify the most appropriate substitution model for tree reconstruction, according to AICc. The best-fitting model was WAG, with r-distributed rate variation across alignment sites (***Gu et al., 1995***). We next performed 300 maximum likelihood tree searches with IQ-TREE (v. 1.6.7; ***Nguyen et al. 2015***) and selected the tree with the highest log-likelihood. For this, we used the default options with the exception of the Nearest Neighbor Interchange tree search, whose thorough mode was activated (flag “-allnni”). The statistical support for the nodes of the best tree was estimated by executing 1,000 nonparametric bootstrap replicates.

#### Species tree reconstruction

To obtain a robust species tree, we collected further molecular sequences of multiple highly phylogenetically informative genes from the species in the study, where possible (Appendix 1— table 2). In particular, our dataset for species tree inference comprised i) amino acid sequences from the two ADK types (which were treated as two distinct genes), ii) 16S and 23S rRNA sequences from the SILVA rRNA database (***Quast et al., 2012***), and iii) amino acid sequences of the 40 protein-coding “PhyEco” marker genes (***Wu et al., 2013***). More precisely, the PhyEco genes are universally present across bacteria and archaea, have been shown to lead to the inference of robust species trees, and exhibit low copy number variation. The study that introduced these 40 genes (***Wu et al., 2013***) had provided FASTA files of bacterial and archaeal amino acid sequences for each PhyEco gene. We filtered these files to extract sequences that belonged to the species in our study, and performed blastp (***Altschul et al., 1997***) searches against the UniProt database to identify sequences from missing species. This process was followed by manual curation to eliminate any false positive hits.

Sequences were aligned with MAFFT as previously described, followed by the removal of phylogenetically uninformative homoplastic sites in each alignment, as identified with Noisy (v. 1.5.12; ***Dress et al. 2008***). We then ran PartitionFinder 2 (***Lanfear et al., 2017***) to determine the optimal partitioning scheme for nucleotide and amino acid sequences. In other words, PartitionFinder 2 identifies groups of genes or proteins (partitions) with similar rates or patterns of evolution, as well as the most appropriate substitution model for them. That model can then be fitted to the entire partition, instead of separately to each gene or protein within the partition. We used the greedy search algorithm (***Lanfear et al., 2012***) of PartitionFinder 2, AICc as the criterion, and RAxML (***Stamatakis, 2014***) for substitution model fitting. The models that were considered were the GTR (for nucleotide sequences) and the WAG, MTREV, DAYHOFF, VT, BLOSUM62, CPREV, RTREV, and MTMAM (for amino acid sequences). The resulting partitioning scheme was used to reconstruct the species phylogeny with IQ-TREE. 20 tree searches were executed, all of which led to the same topology (Appendix 1—figure 4). As previously, the statistical support was estimated via 1,000 bootstrap replicates.

For time-calibration of the phylogeny, we used BEAST (v. 2.5.2; ***Bouckaert et al. 2014***) and a lognormal relaxed clock (***Drummond et al., 2006***), keeping the topology fixed to that inferred by IQ-TREE. We used the same partitioning scheme as before, but replaced any substitution models that were not available in BEAST 2 with the general WAG model. To obtain a starting tree with ultrametric branch lengths, we used the chronos function in the ape R package (v. 5.2; ***Paradis and Schliep 2019***), which is based on penalised likelihood (***Paradis, 2013***). Two independent BEAST 2 runs were executed for 450 million generations, with posterior samples being obtained every thousand generations, after discarding samples from the first 112.5 million generations as burn-in. To ensure that the two runs converged to a common posterior distribution, we calculated the effective sample size and the potential scale reduction factor for each model parameter (***Gelman and Rubin 1992***; ***Brooks and Gelman 1998***; Appendix 1—figure 5). Our convergence criteria were i) an effective sample size of at least 200 and ii) a potential scale reduction factor value below 1.1. The same convergence diagnostics were performed for the estimated substitution rate of each tree branch (Appendix 1—figure 6). We summarised the posterior distribution of trees by calculating the median age estimate for each node. As we had not fixed the age for any node, the branch lengths of the resulting tree were in units of substitutions per site. To convert them to units of time, we set the root to a depth of 3,700 mya, according to current estimates of the date of the origin of life on Earth (***Pearce et al., 2018***). Substitution rates were also divided by 3,700.

### Molecular dynamics simulations

#### Simulation protocol

Molecular dynamics simulations were performed using the Desmond software and its graphical interface, Maestro (Version 10.6.013, Release 2016-2). Our protocol was based on Desmond’s molecular dynamics simulation tutorial, distributed with the software suite.

To prepare each ADK for the simulations, we first capped the N- and C-termini and added missing hydrogens. To remove close contacts or clashes between atoms, we relaxed the structure into a local energy minimum, in vacuum (i.e., in the absence of solvent), using the OPLS 2005 force field (***Banks et al., 2005***). The minimized protein was solvated in a periodic orthorhombic simulation box with Simple Point Charge water molecules (***Berendsen et al., 1981***). Na^+^ and Cl^−^ ions were added to neutralize the charge of the protein as appropriate, and to generate a NaCl salt concentration of 0.15 M. The resulting system was relaxed using Desmond’s standard protocol.

We next performed ten independent simulations (with randomised initial atomic velocities) per ADK. These were performed under the NPT ensemble, i.e., with temperature and pressure being maintained constant using the Nosé-Hoover chain thermostat (***Martyna et al., 1992***) and the Martyna-Tobias-Klein barostat (***Martyna et al., 1994***) respectively. We varied the temperature of the simulations by the thermal group: 280 K (6.85°C) for psychrophiles, 300 K (26.85°C) for mesophiles, 330 K (56.85°C) for thermophiles, and 355 K (81.85°C) for hyperthermophiles. The pressure was kept constant at the default value of 1.01325 bar (the standard atmospheric pressure) using isotropic coupling and a relaxation time of 2 ps. The cutoff radius for van der Waals and short range electrostatic interactions was set to 9 Å. The integration time step was left at the default value of 2 fs, with snapshots of the system being recorded every 500 ps until each simulation was 200 ns long.

#### Trajectory analysis

We analyzed the ten replicate molecular dynamics trajectories per ADK by first calculating the root-mean-square deviation (RMSD) matrix across all recorded snapshots (conformations) of the protein. The RMSD between two conformations of the same protein, **a** and **b**, is defined as

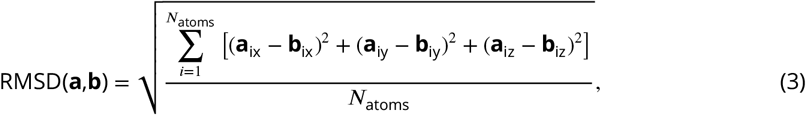

where x, y, and z are the axes of the three-dimensional space. As the entire protein moves in space during the course of a molecular dynamics simulation, an objective comparison of two conformations requires finding the superposition between them that minimizes the RMSD.

It is worth stressing that ADK structures contain a large number of amino acids that participate in neither helices nor *β*-strands. Such amino acids are usually highly flexible and their motions are largely random. Therefore, to reduce the influence of highly flexible amino acids on the identification of protein conformations, we excluded such amino acids from the calculation of the RMSD matrix. For this, we converted the trajectories from the Desmond format to DCD format with the VMD program (***Humphrey et al., 1996***). VMD was also used to generate a Protein Structure File. These two files were then analysed with Carma (***Glykos, 2006***). Carma can automate the secondary structure analysis of molecular dynamics trajectories by calling the STRIDE secondary structure assignment tool (***Frishman and Argos, 1995***) and analysing its output. This allowed us to identify the amino acids of each ADK that participated in neither helices nor *β*-strands for at least 80% of the combined duration of the ten replicate trajectories. Such amino acids were excluded from the superposition and the RMSD calculation.

The resulting RMSD matrix was clustered using the Partitioning Around Medoids algorithm (***Reynolds et al., 2006***), as implemented in the cluster R package (***Maechler et al., 2017***). The only requirement of this algorithm is the number of clusters to detect, whereas the optimal number of clusters can be determined by comparing the mean silhouette score (***Rousseeuw, 1987***) of different clusterings. In our case, we set the initial number of clusters to all possible values between 2 and 20, and selected the clustering with the highest mean silhouette score. Finally, we used Carma to obtain the average (representative) conformation for each cluster.

### The relationship between median mutational effects and temperature across species

To check for a systematic effect of temperature on 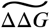 values of different species, we fitted a series of models with MCMCglmm. The response variable was 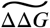, whereas possible predictors (other than an intercept) included: i) the temperature at which molecular dynamics simulations were performed, ii) the ADK type (ancestral or archaeal), and iii) the length of the LID. All possible combinations of these predictors were specified in different models, including an empty model that only contained an intercept. A phylogenetic variant of each model was also fitted, using the species phylogeny that we inferred. In that case, species identity was treated as a random effect on the intercept. An estimate of uncertainty for each 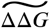 value was obtained through bootstrapping and was integrated into the analysis. We used the default prior for the fixed effects, and a relatively uninformative inverse-r prior for the random effect and residual variances. For each model, two replicate MCMC chains were executed for three million generations, with samples being obtained every thousand generations. We discarded samples from the first three hundred thousand generations as burn-in and ensured that the two chains per model had suZciently converged on identical posterior distributions by examining the effective sample size for each parameter and the potential scale reduction factor, as described earlier. For identifying the most appropriate model, we first rejected models that had at least one coefficient whose 95% Highest Posterior Density (HPD) interval included zero. From the remaining models, we chose the model with the lowest Deviance Information Criterion (***Spiegelhalter et al. 2002***; DIC) value.

We also estimated the phylogenetic heritability of 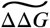 to understand if closely related species have more similar values than species chosen at random. This involved fitting a phylogenetic regression of 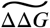 using MCMCglmm with only an intercept as a predictor. This model was fitted separately to each ADK type by executing two replicate chains for ten million generations, with samples being obtained every thousand generations after the first million generations. The phylogenetic heritabilities of 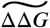 for the two ADK types were then estimated as the ratio of the variance explained by the random effect (species identity corrected for phylogeny) over the sum of the random effect variance and the residual variance.

To visualise the evolution of 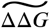 for each ADK type on the species phylogeny (Figure 4B,C), we estimated ancestral states by fitting the stable model of trait evolution (***Elliot and Mooers, 2014***). For this, four MCMC chains were executed for a hundred million generations, sampling every hundred generations after a burn-in phase of 25 million generations. Convergence of the four chains was examined as previously described.

### Estimation of the relationship between substitution rate and temperature

We fitted variants of Equation (1) with MCMCglmm, with and without a phylogenetic correction on the intercept by executing two MCMC chains per model for a million generations, with posterior samples being recorded every thousand generations after the first hundred thousand. Because cell volume estimates were available as ranges for most species, we fitted models that included cell volume 20 times. Each time, the volume of each species was sampled from a uniform distribution with bounds that corresponded to the minimum and maximum volume calculations.

## Acknowledgments

We thank Hira Tanvir for providing some of the cell volume data that were used in this study, and Mark E. Ritchie, Andrew Hirst, Guy Woodward, Vassiliki Koufopanou, Bhavin S. Khatri, Vincent Savolainen, and the Michael J. E. Sternberg Lab for useful discussions and comments. We are also particularly grateful to the developers and maintainers of the STRUM web service, as a major component of this study depended on it. Finally, we thank Imperial College London’s High Performance Computing service (doi:10.14469/hpc/2232) for access to the necessary computational resources for performing the molecular dynamics simulations.

## Additional files

### Data availability

The data and source code that support the conclusions of this study are available at https://doi.org/10.6084/m9.figshare.12635837.v1 and https://github.com/dgkontopoulos/Kontopoulos_et_al_mutations_vs_temperature_2020 respectively.

## APPENDIX 1

## I. The Gibbs-Helmholtz equation

If all environmental variables other than temperature (e.g., pH, salt concentration) are kept constant, the relationship between protein stability (Δ*G*) and temperature (*T*) for proteins unfolding according to the two-state model can be described by the Gibbs-Helmholtz equation (***Privalov, 1990***):

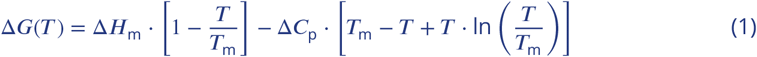

Here, *T*_m_ (the “melting temperature”, in K units) is the temperature at which Δ*G* = 0 due to heat denaturation. At *T*_m_, the protein population is equally distributed between the two states (folded and unfolded). Δ*H*_m_ (kcal · mol^−1^) is the difference in enthalpy between the folded and unfolded states at *T*_m_. Δ*C*_p_ (kcal · mol^−1^ · K^−1^) is the difference in heat capacity between the two states, which is assumed to be independent of temperature.

**Appendix 1—figure 1.**
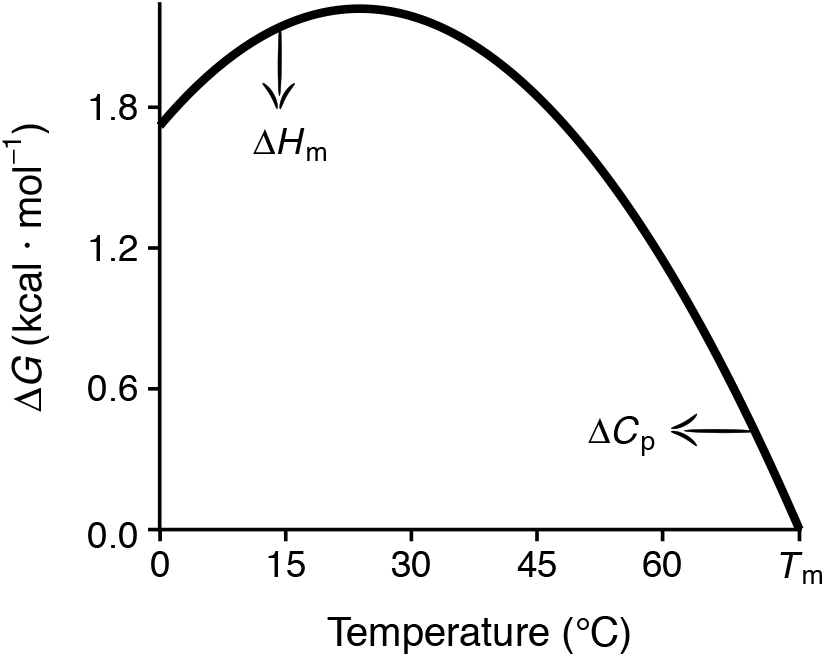
An example of a protein stability curve according to the Gibbs-Helmholtz equation. The temperature is shown in °C for clarity but in the equation it is in K units. Note that Δ*H*_m_, which mainly controls the maximum height of the stability curve, is normalised at *T* = *T*_m_.

## II. List of analysed prokaryotic ADKs

**Appendix 1—table 1.**
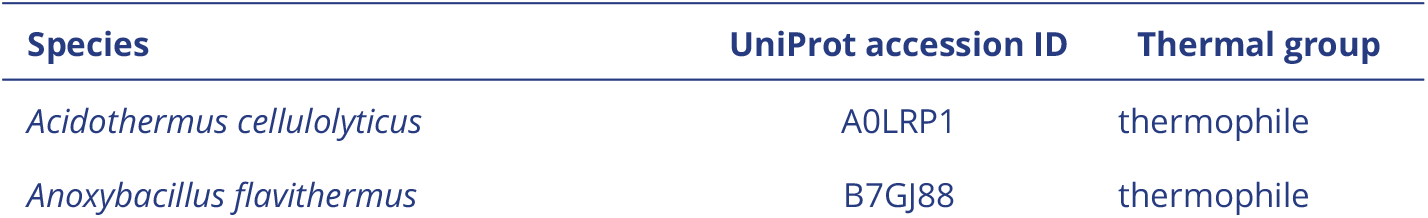
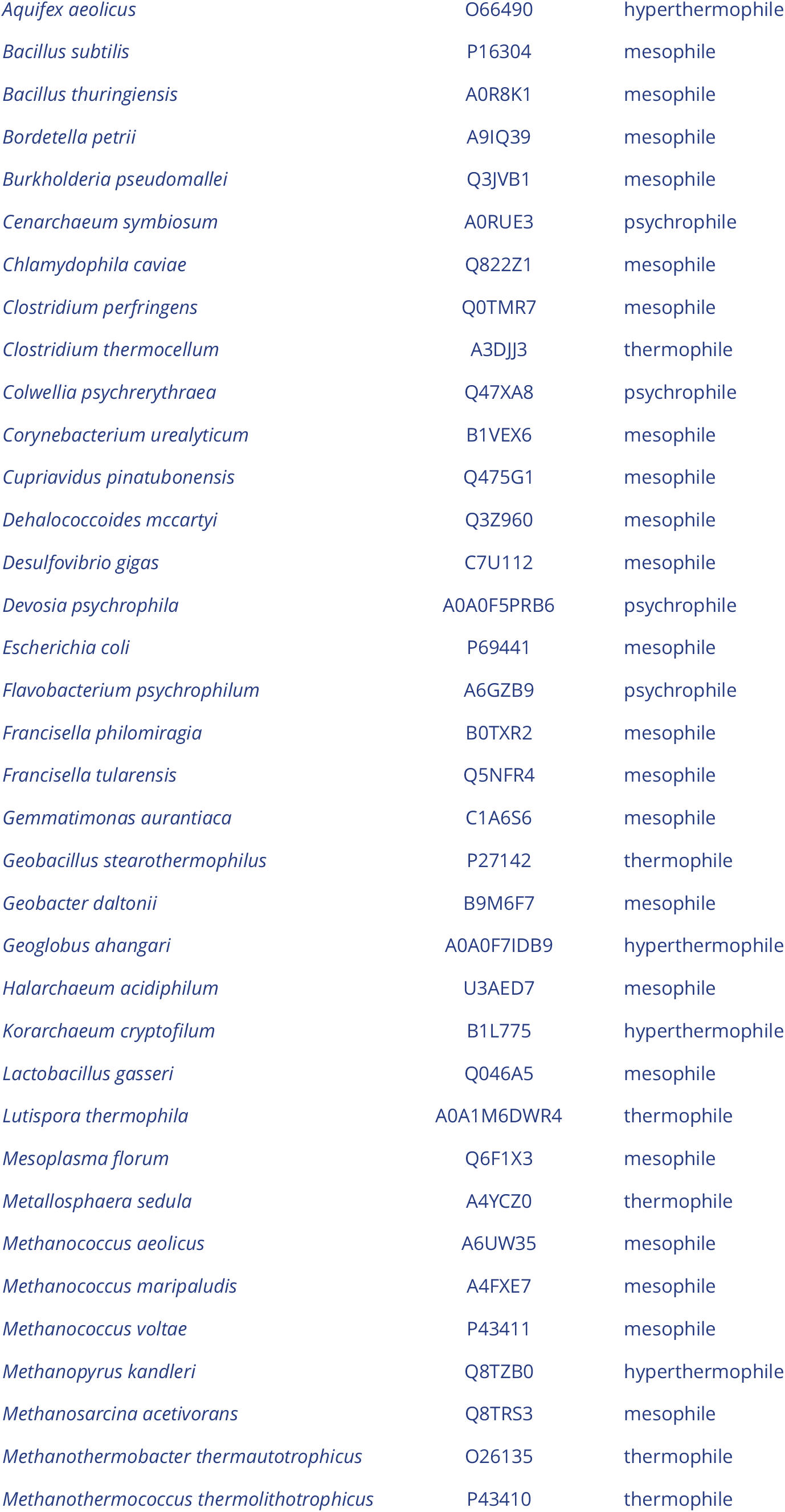
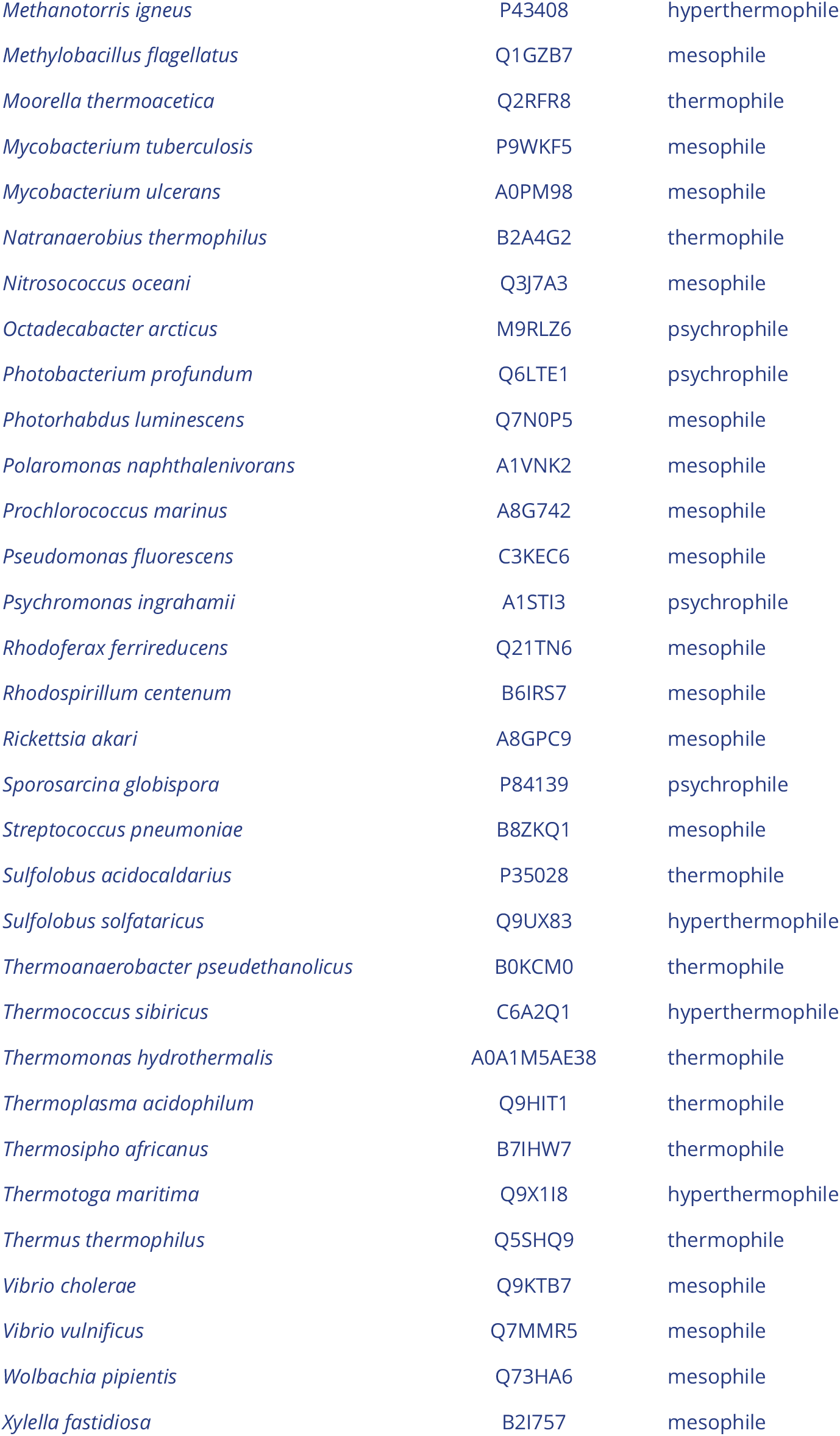
Prokaryotic ADKs that were included in this study.

## III. Homology modelling of prokaryotic ADKs

**Appendix 1—figure 2.**
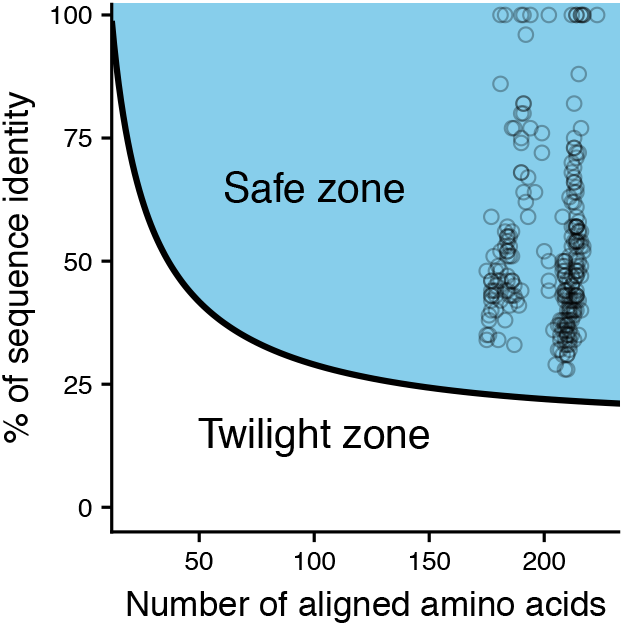
Percentage of sequence identity versus alignment length for all template sequences of this study. Note that many of our homology models were built using multiple templates and, therefore, multiple data points can correspond to the same homology model. Data points at 100% sequence identity indicate ADKs whose structures were available from the PDB, and for which homology modelling was done only to add any missing amino acids or not at all.

**Appendix 1—figure 3.**
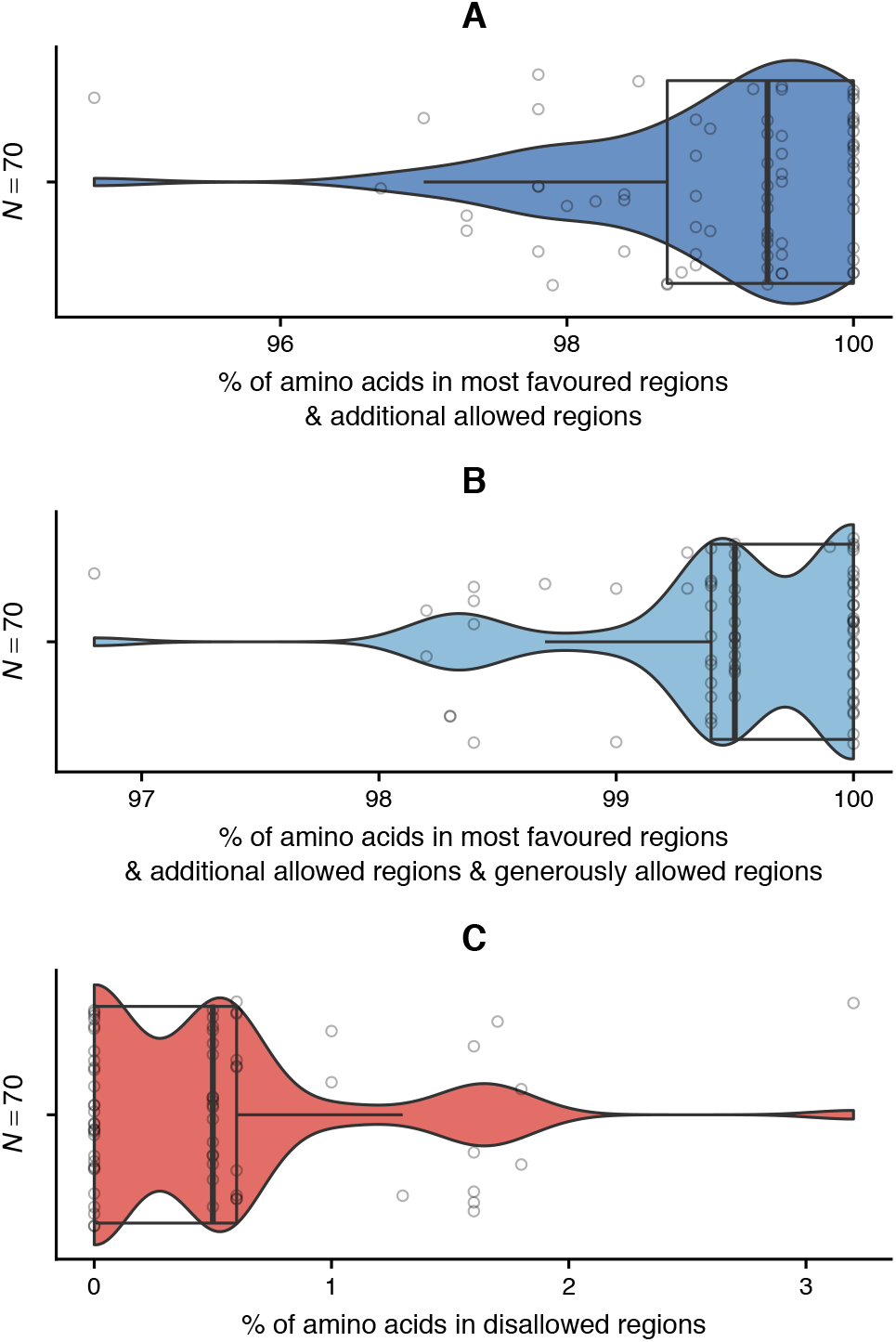
Distribution of amino acids across the regions of the Ramachandran plot for each ADK structure of our dataset. Boxplot edges represent the first and third quartiles, whereas the solid line stands for the median. Whiskers extend up to the most remote data point within 1.5 interquartile ranges from each boxplot edge. Given that all structures had >94% of their amino acids in the most favoured and additional allowed regions, and at worst just over 3% of amino acids in disallowed regions, their quality can be considered sufficiently high.

## IV. Species tree reconstruction

Appendix 1—table 2 lists the genes from which we reconstructed the species tree. The best partitioning scheme is shown in Appendix 1—table 3. It is worth noting that the two ADK types were assigned to two different partitions and models. The resulting species phylogeny is shown in Appendix 1—figure 4. Finally, Appendix 1—figures 5 and 6 show convergence diagnostics for the time-calibration of the phylogeny with BEAST 2.

**Appendix 1—table 2.** Molecular sequences used for species tree inference.

| ID | Description | Type |
| --- | --- | --- |
| 16S | Small subunit rRNA | Nucleotide |
| 23S | Large subunit rRNA | Nucleotide |
| ADK (ancestral type) | Adenylate kinase | Amino acid |
| ADK (archaeal type) | Adenylate kinase | Amino acid |
| PhyEco_BA_01 | Ribosomal protein S2 | Amino acid |
| PhyEco_BA_02 | Ribosomal protein S10 | Amino acid |
| PhyEco_BA_03 | Ribosomal protein L1 | Amino acid |
| PhyEco_BA_04 | Translation elongation factor EF-2 | Amino acid |
| PhyEco_BA_05 | Translation initiation factor IF-2 | Amino acid |
| PhyEco_BA_06 | Metalloendopeptidase | Amino acid |
| PhyEco_BA_07 | Ribosomal protein L22 | Amino acid |
| PhyEco_BA_08 | Ffh signal recognition particle protein | Amino acid |
| PhyEco_BA_09 | Ribosomal protein L4/L1e | Amino acid |
| PhyEco_BA_10 | Ribosomal protein L2 | Amino acid |
| PhyEco_BA_11 | Ribosomal protein S9 | Amino acid |
| PhyEco_BA_12 | Ribosomal protein L3 | Amino acid |
| PhyEco_BA_13 | Phenylalanyl-tRNA synthetase $\beta$ subunit | Amino acid |
| PhyEco_BA_14 | Ribosomal protein L14b/L23e | Amino acid |
| PhyEco_BA_15 | Ribosomal protein S5 | Amino acid |
| PhyEco_BA_16 | Ribosomal protein S19 | Amino acid |
| PhyEco_BA_17 | Ribosomal protein S7 | Amino acid |
| PhyEco_BA_18 | Ribosomal protein L16/L10E | Amino acid |
| PhyEco_BA_19 | Ribosomal protein S13 | Amino acid |
| PhyEco_BA_20 | Phenylalanyl-tRNA synthetase $\alpha$ subunit | Amino acid |
| PhyEco_BA_21 | Ribosomal protein L15 | Amino acid |
| PhyEco_BA_22 | Ribosomal protein L25/L23 | Amino acid |
| PhyEco_BA_23 | Ribosomal protein L6 | Amino acid |
| PhyEco_BA_24 | Ribosomal protein L11 | Amino acid |
| PhyEco_BA_25 | Ribosomal protein L5 | Amino acid |
| PhyEco_BA_26 | Ribosomal protein S12/S23 | Amino acid |
| PhyEco_BA_27 | Ribosomal protein L29 | Amino acid |
| PhyEco_BA_28 | Ribosomal protein S3 | Amino acid |
| PhyEco_BA_29 | Ribosomal protein S11 | Amino acid |
| PhyEco_BA_30 | Ribosomal protein L10 | Amino acid |
| PhyEco_BA_31 | Ribosomal protein S8 | Amino acid |
| PhyEco_BA_32 | tRNA pseudouridine synthase B | Amino acid |
| PhyEco_BA_33 | Ribosomal protein L18P/L5E | Amino acid |
| PhyEco_BA_34 | Ribosomal protein S15P/S13e | Amino acid |
| PhyEco_BA_35 | Porphobilinogen deaminase | Amino acid |
| PhyEco_BA_36 | Ribosomal protein S17 | Amino acid |
| PhyEco_BA_37 | Ribosomal protein L13 | Amino acid |
| PhyEco_BA_38 | Phosphoribosylformylglycinamide cyclo-ligase | Amino acid |
| PhyEco_BA_39 | Ribonuclease HII | Amino acid |
| PhyEco_BA_40 | Ribosomal protein L24 | Amino acid |

**Appendix 1—table 3.**
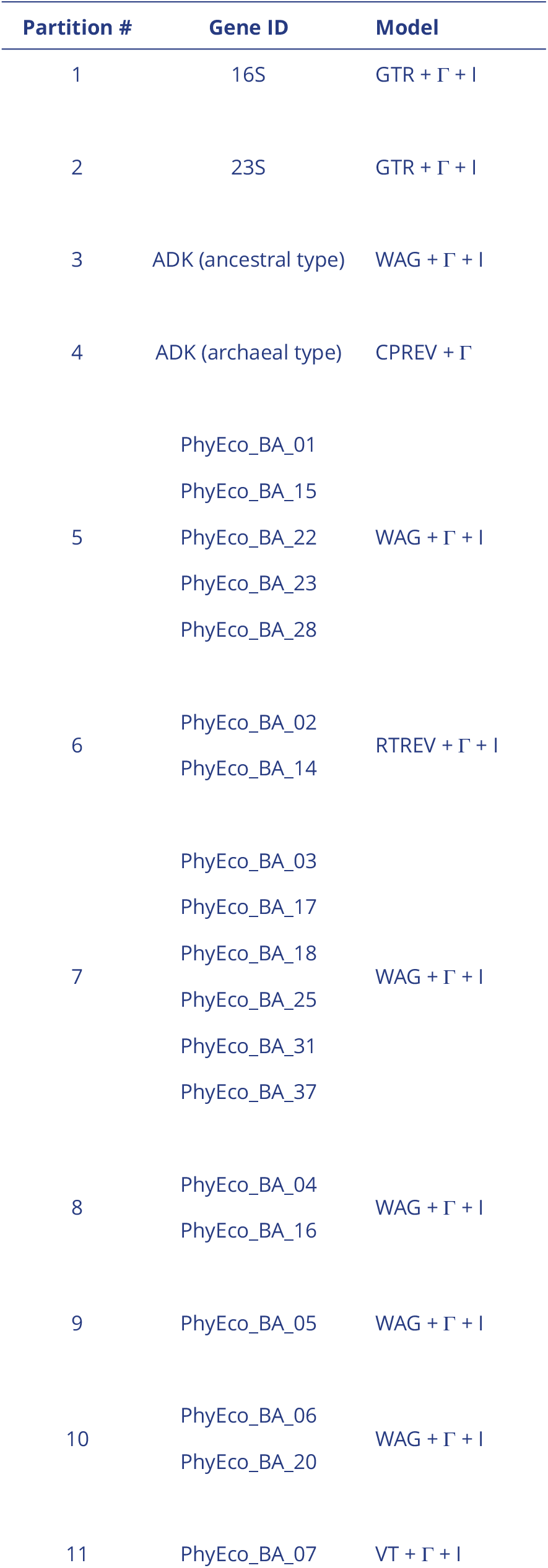
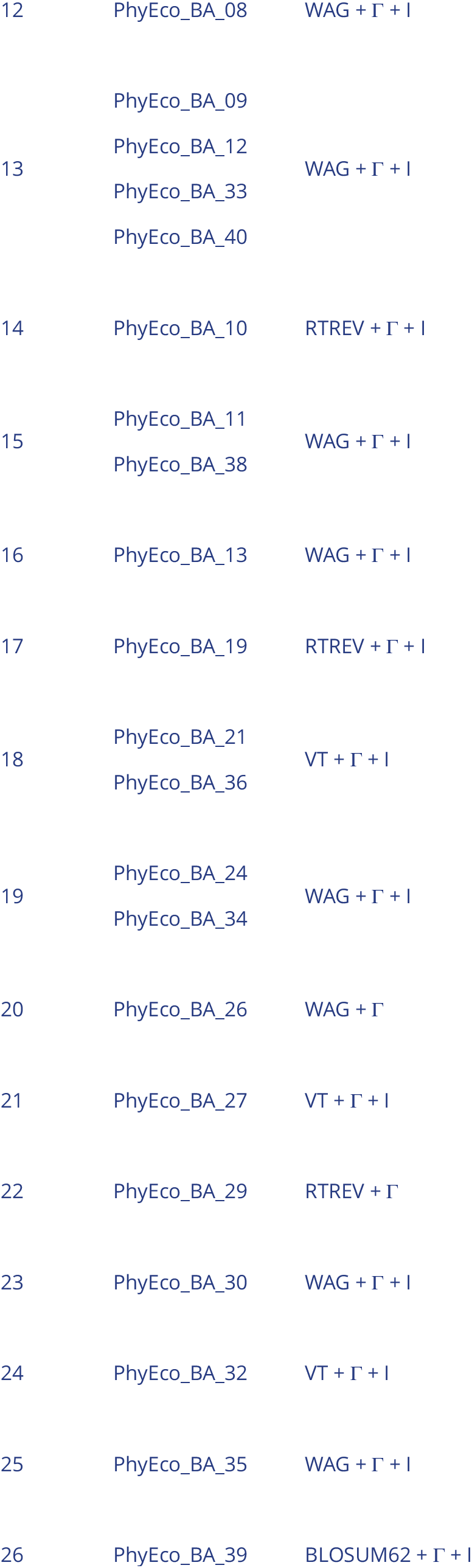
Optimal partitioning scheme according to AICc.

**Appendix 1—figure 4.**
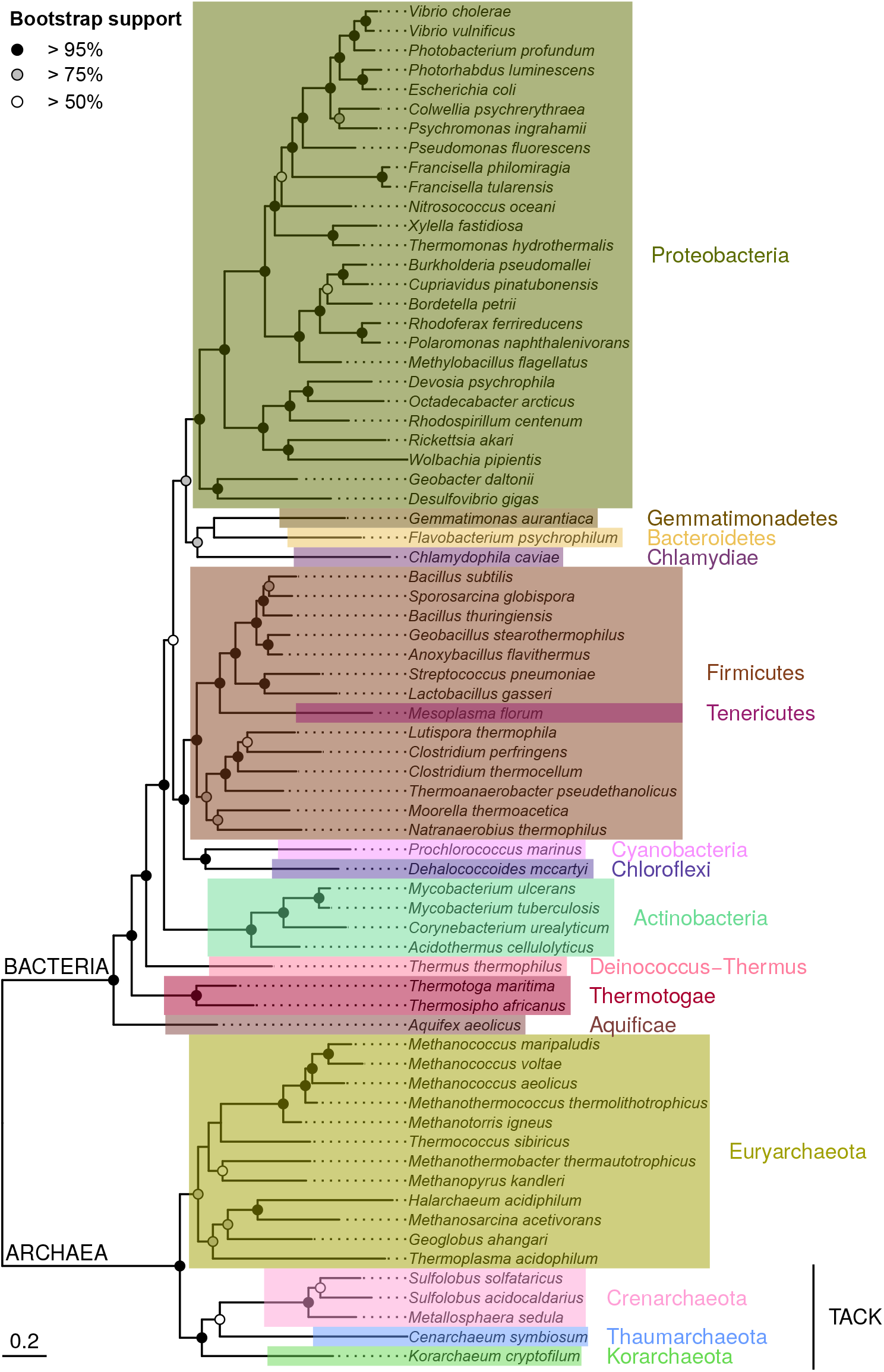
The maximum likelihood phylogeny of the species in our dataset of ADKs, as inferred with IQ-TREE. Each phylum is shown in a different colour. Additionally, the phyla that belong to the TACK superphylum are explicitly marked. In contrast to the ADK gene tree, most nodes of the species tree have sufficiently high statistical support (black and grey circles).

**Appendix 1—figure 5.**
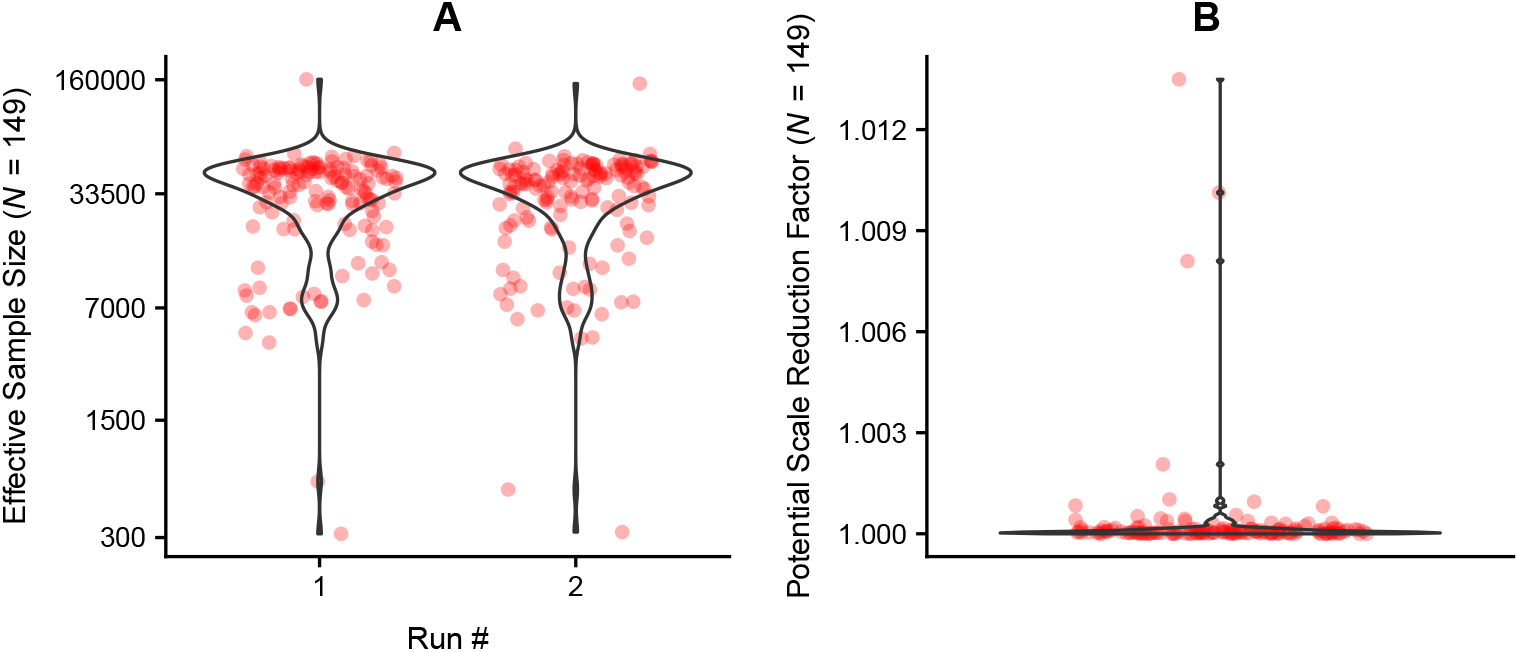
Effective sample size and potential scale reduction factor values for each parameter of the evolutionary model fitted with BEAST 2.

**Appendix 1—figure 6.**
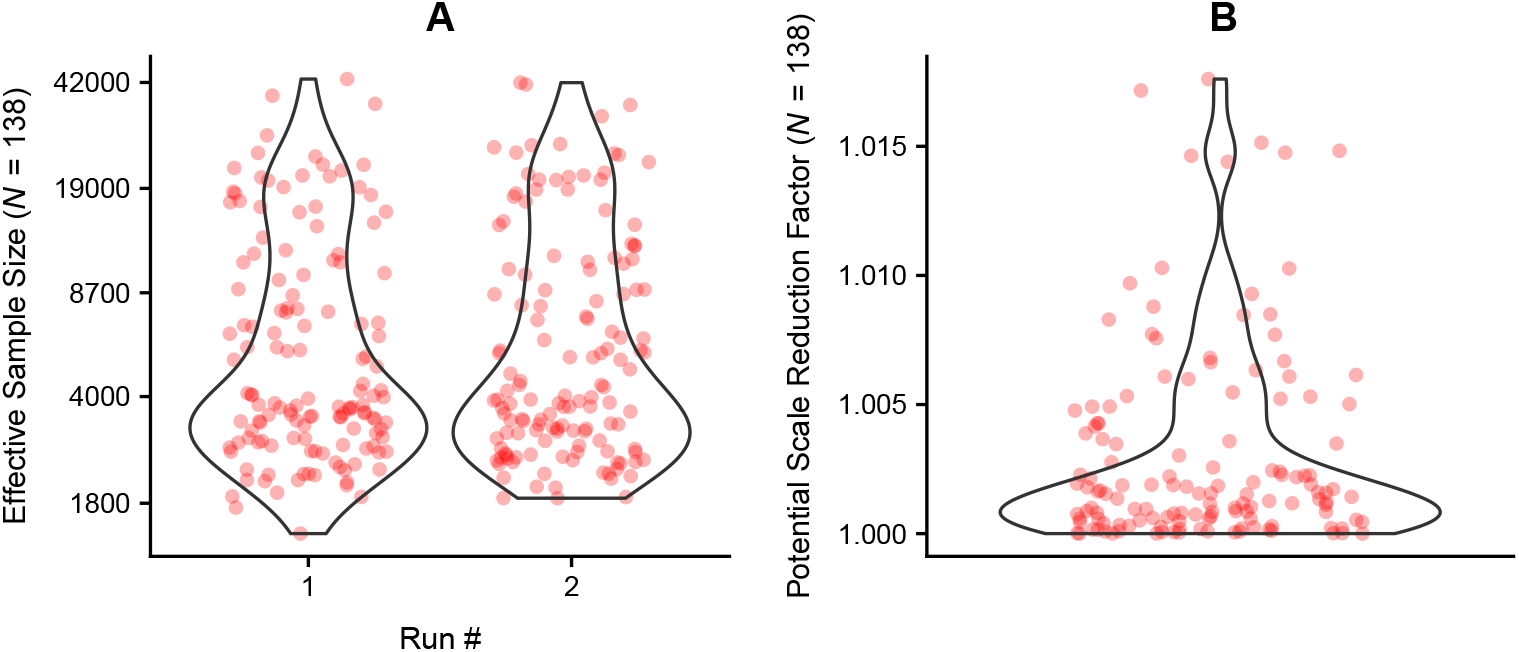
Effective sample size and potential scale reduction factor values for each branch-specific substitution rate, as estimated with BEAST 2.

## V. Analysis of molecular dynamics trajectories

**Appendix 1—figure 7.**
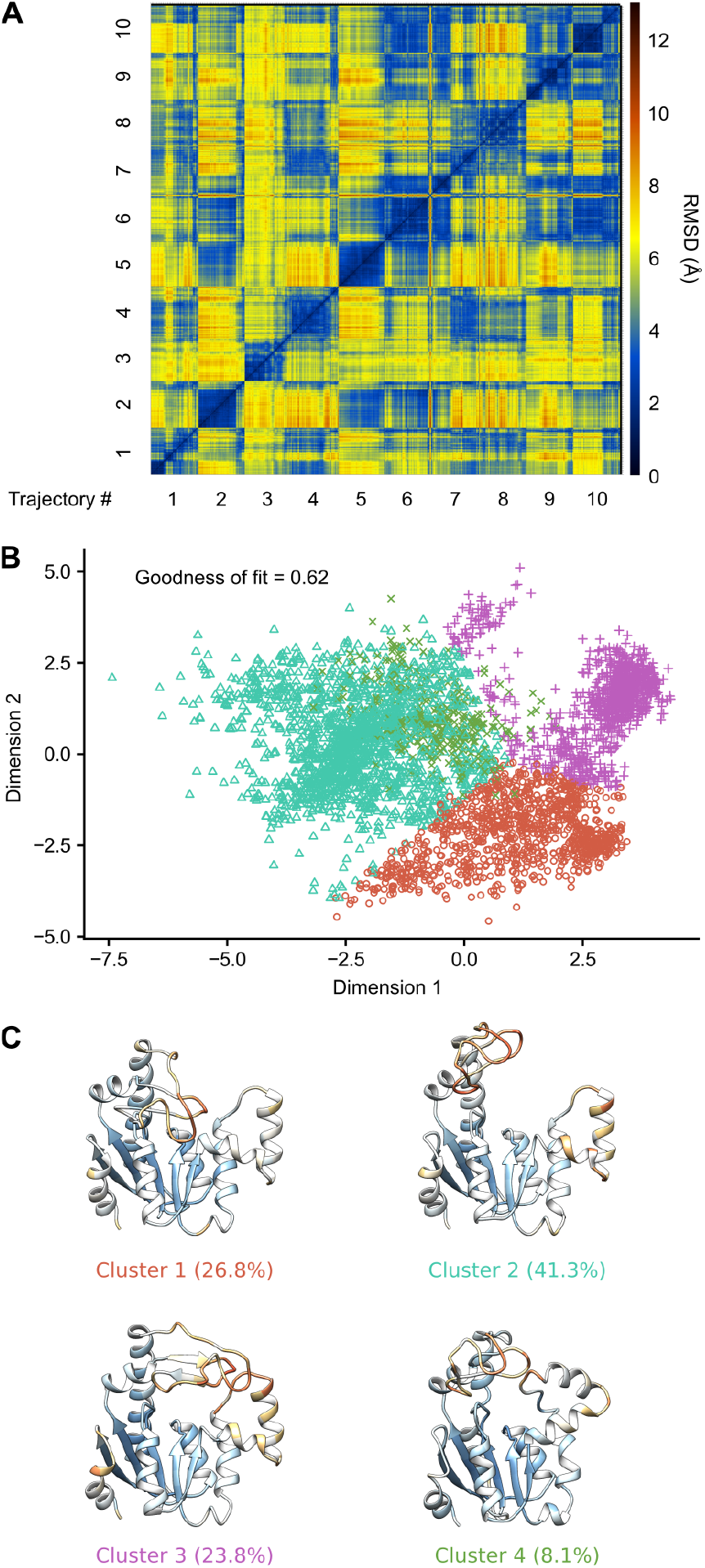
Illustration of the procedure for obtaining representative ADK conformations, based on the molecular dynamics trajectories of *Francisella philomiragia*. A: The RMSD matrix among all snapshots is calculated after excluding highly flexible amino acids. B: Conformations are grouped into major structural clusters, visualised here using multidimensional scaling. Note that the reported goodness of fit refers to the ability of two dimensions to properly reflect the RMSD values among data points, and is not a measure of clustering quality. C: From each cluster, an average structure is produced. The flexibility of amino acids for each conformation is shown on a colour scale that ranges from dark orange (highly flexible) to white to dark blue (least flexible). The percentages show the proportion of protein snapshots that belong to each conformation.

## VI. Investigation of biases in the estimates of mutational effects

**Appendix 1—figure 8.**
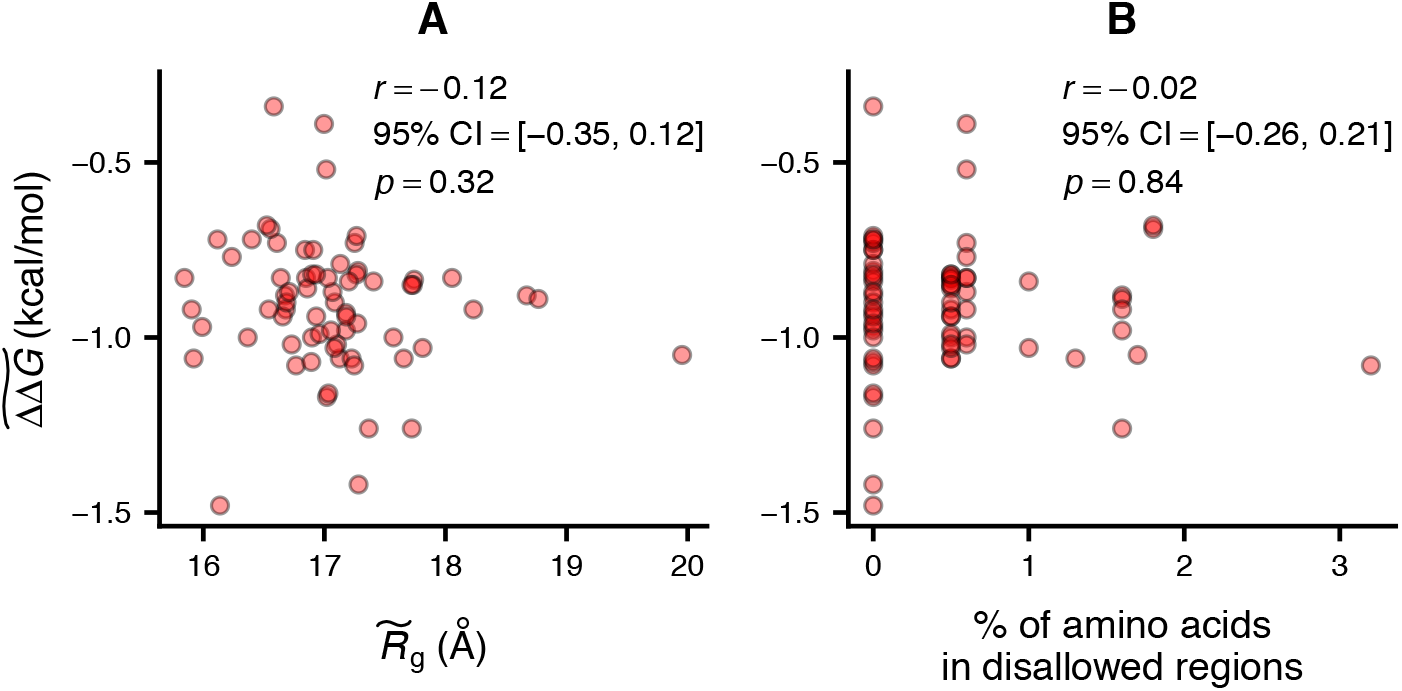
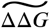 estimates were not found to be influenced by either the radius of gyration (panel A) or the quality of the homology model (panel B).

## VII. The inter- and intraspecific relationships between median mutational effects and temperature

Appendix 1—table 4 lists the models that were fitted to characterise the relationship between 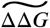 and temperature across species. Models with at least one coefficient whose 95% HPD interval included zero are not shown.

**Appendix 1—table 4.** Candidate models for the interspecific relationship between 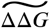 and temperature. The model with the lowest DIC is highlighted.

| Model | Phylogenetic correction | Mean DIC |
| --- | --- | --- |
| $\widetilde{\Delta\Delta G} \sim \text{Intercept}$ | ✓ | -27.1691 |
| $\widetilde{\Delta\Delta G} \sim \text{Intercept} + \text{Temperature}$ | ✓ | -32.4251 |
| $\widetilde{\Delta\Delta G} \sim \text{Intercept} + \text{ADK type}$ | ✓ | -30.8497 |
| $\widetilde{\Delta\Delta G} \sim \text{Intercept} + \text{Temperature} + \text{ADK type}$ | ✓ | -35.3310 |
| $\widetilde{\Delta\Delta G} \sim \text{Intercept}$ | ✗ | -28.4569 |
| $\widetilde{\Delta\Delta G} \sim \text{Intercept} + \text{LID length}$ | ✗ | -33.9572 |
| $\widetilde{\Delta\Delta G} \sim \text{Intercept} + \text{ADK type}$ | ✗ | -53.9365 |
| $\widetilde{\Delta\Delta G} \sim \text{Intercept} + \text{Temperature} + \text{ADK type}$ | ✗ | -63.9542 |

To estimate the species-specific relationships between median ΔΔ*G* and temperature, we fitted six mathematical equations to the data (Appendix 1—figure 9). The most appropriate shape for each species (Figure 5) was determined using AICc.

**Appendix 1—figure 9.**
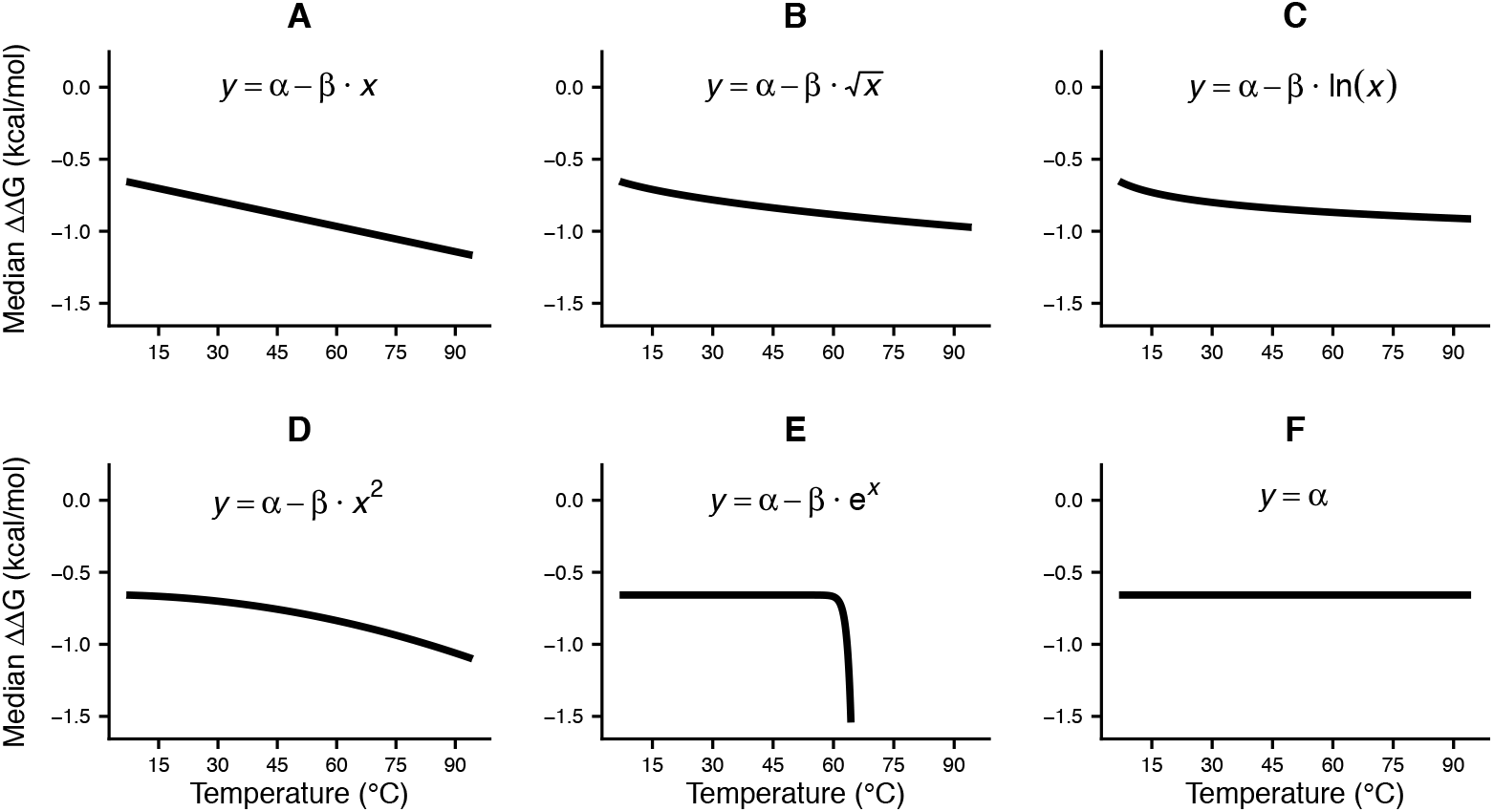
The six mathematical models that we fitted to the dataset of median ΔΔ*G* versus temperature for each ADK. These models capture a wide range of shapes, from a linear decrease to nonlinear changes or even a constant value across temperatures.

Due to the variation in the mathematical form of the fitted equations, it was not possible to directly compare the relationships on the basis of model coefficients. For an objective comparison, we instead executed PCA on a matrix of predicted median ΔΔ*G* values (for each ADK) at temperatures from 6.85°C to 94.35°C with a step of 0.05°C. This showed that species-specific relationships can be adequately described using two principal components, on which we also projected the phylogeny (Appendix 1—figure 10). The first component captures 95.4% of the variance and stands for the average robustness to mutations across all temperatures. The second principal component represents the temperature sensitivity of the median ΔΔ*G*. The values of internal nodes (except for the node corresponding to LUCA) were estimated using the stable model of trait evolution (***Elliot and Mooers, 2014***). More precisely, we executed four MCMC chains for 1.5 billion generations, sampling every 2,000 generations after the first 375 million generations.

**Appendix 1—figure 10.**
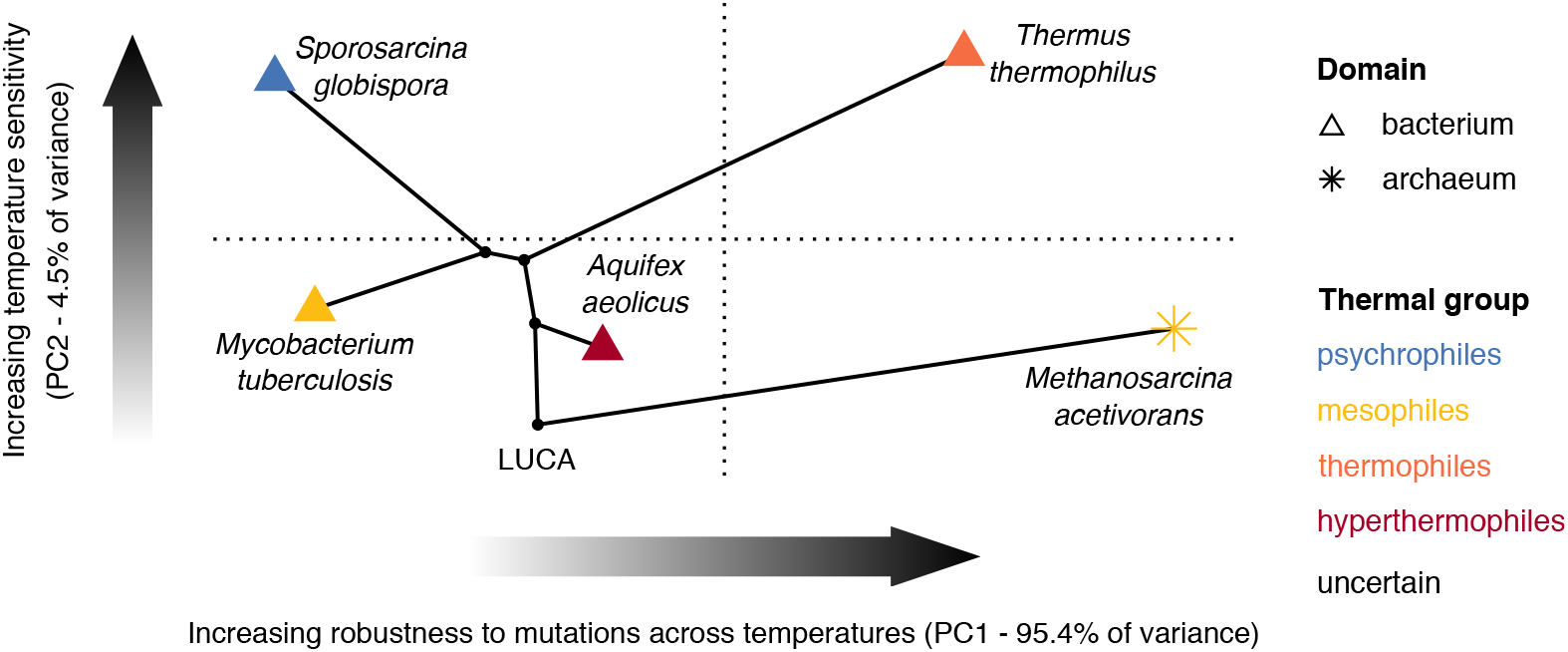
Projection of the relationships between median ΔΔ*G* and temperature to the first two principal components. Most of the variance (PC1 axis) corresponds to shifts in mutational robustness (i.e., changes along the vertical axis of plots of Figure 5). A much smaller amount of variance (PC2 axis) is due to variation in the temperature sensitivity of median ΔΔ*G* among ADKs of different species.

It is worth noting that the ADK of the LUCA had the lowest temperature sensitivity across the six species, suggesting a likely evolutionary trend towards increasing temperature sensitvity. However, given that i) the ADK gene is under strong selection and that ii) it evolves rapidly and in a convergent manner across species (Figure 3A), such a trend in temperature sensitivity could be an artefact of the ancestral reconstruction of the LUCA sequence. To understand whether this trend is real, future studies could perform a similar analysis for genes that are characterised by non-convergent and less rapid evolution.

## VIII. The ADK of the Last Universal Common Ancestor

## VIII.I Estimated sequence and 3D structure

To reconstruct the ADK sequence of the LUCA, we submitted to the FASTML server (***Ashkenazy et al., 2012***) the alignment of ancestral-type ADKs, the species tree, and the most appropriate model of sequence substitution (WAG; Appendix 1—table 3). The resulting sequence is as follows:

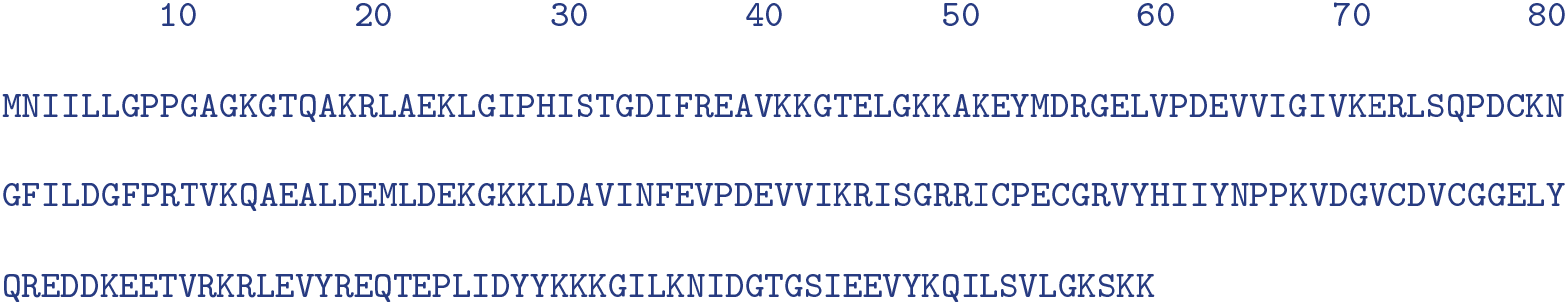

Its homology model that we obtained via Phyre2 is shown in Appendix 1—figure 11.

**Appendix 1—figure 11.**
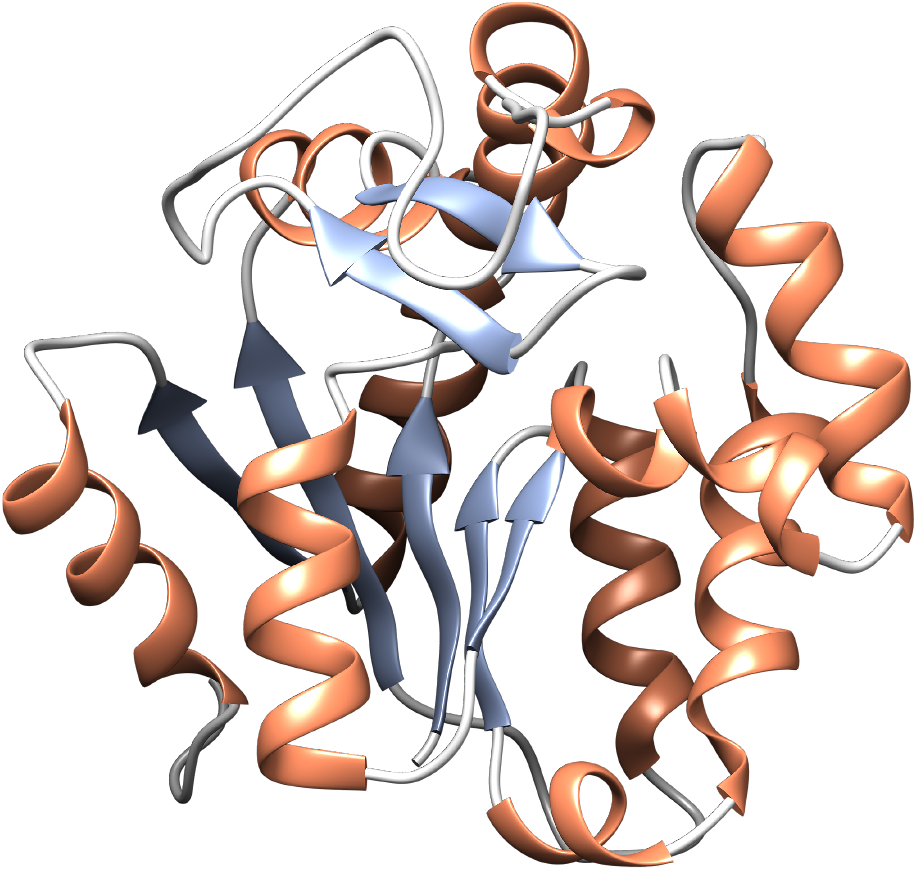
The inferred ADK structure of the LUCA. Helices are shown in orange, whereas *β*-strands are in blue. The LID domain is long, similarly to that of most ancestral-type ADKs.

## VIII.II Estimation of the temperature of optimal catalysis (*T_opt_*)

In microbes, the temperature of optimal catalysis of a given enzyme (*T*_opt_) has been found to correlate with the temperature of maximum growth rate (*T*_pk_; ***Engqvist 2018***; ***Li et al. 2019***). The strength of this correlation varies across enzymes, possibly because the activity of certain enzymes (e.g., ADKs; ***Couñago and Shamoo 2005***; ***Couñago et al. 2006***) is more tightly linked to organismal fitness than others. Thus, estimating *T*_opt_ for the ADK of the LUCA should provide a rough understanding of the environmental conditions in which LUCA emerged.

***Li et al.*** (***2019***) showed that the *T*_opt_ of an enzyme can be relatively accurately predicted using a random forest model. The inputs of the model are i) amino acid frequencies for the enzyme of choice and ii) the organism’s growth rate *T*_pk_. The model can also be trained without *T*_pk_ data, but at a cost of slightly worse performance. To this end, we used the code and training dataset of ***Li et al.*** (***2019***) to build a random forest model that would predict *T*_opt_ from amino acid frequencies alone. The resulting model achieved an *R*^2^ of 0.9193 (RMSE = 5.328) against its training dataset. We next used this model to infer the likely *T*_opt_ value of the ADK sequence of the LUCA (Appendix 1—figure 12).

**Appendix 1—figure 12.**
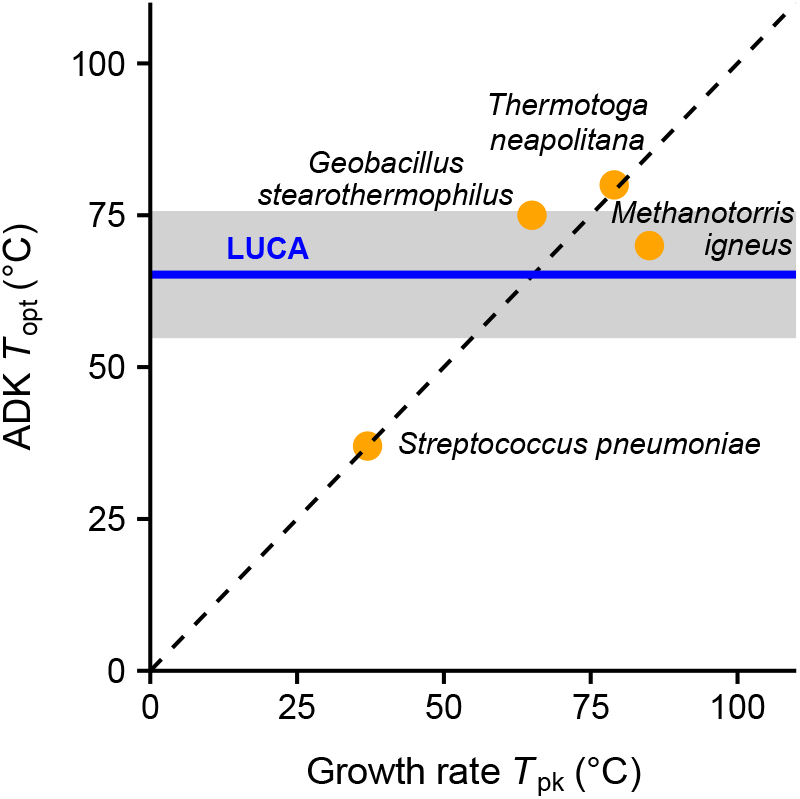
*T*_opt_ values of prokaryotic ADKs against the temperature of maximum growth rate (*T*_pk_). The dashed line is the one-to-one line, the estimated *T*_opt_ of the LUCA is shown in blue, whereas the 95% prediction interval is shown in grey. Even though the characteristics of the thermal environment of the LUCA remain under debate, the close relationship between *T*_opt_ and *T*_pk_ in four other species suggests that LUCA was likely a moderate thermophile. The data points were taken from ***Couñago and Shamoo*** (***2005***) and ***Li et al.*** (***2019***).

## IX. Estimation of cell volume from cell dimension measurements

We estimated the cell volume (*V*) of spherical microbes (Appendix 1—figure 13A) as

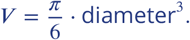

For microbes whose shape was reported as similar to a disc, a filament, or a cylinder (Appendix 1—figure 13B), we used the following equation:

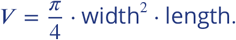

Last, for rod-shaped microbes (Appendix 1—figure 13C), we estimated their cell volume as the sum of the volumes of a cylinder and two half-spheres:

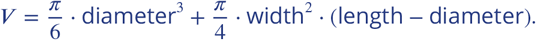

**Appendix 1—figure 13.**
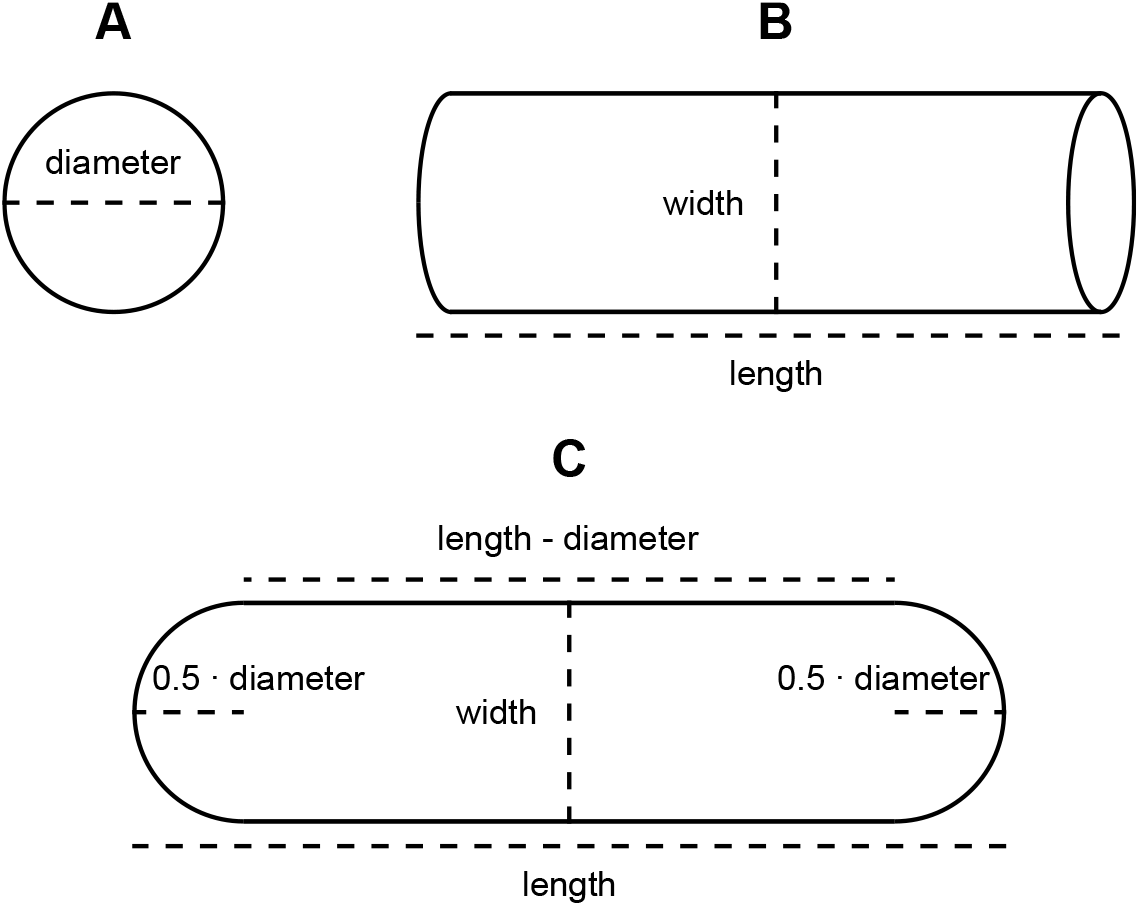
The three-dimensional geometrical shapes based on which we estimated microbial cell volumes in this study. Note that in panel C, the diameter of the two half-spheres is equal to the width of the cell.

## X. Estimation of prokaryotic generation times as a function of temperature and/or cell volume

## X.I Temperature

The relationship between population growth rate and temperature in ectotherms has the shape of a unimodal curve (the thermal performance curve; ***Angilletta 2009***; Appendix 1—Figure 14A). As previously mentioned, the temperature at which the curve peaks is defined as *T*_pk_, whereas the population growth rate at *T*_pk_ is *B*_pk_. *T*_pk_ tends to be slightly above the mean environmental temperature (***Martin and Huey, 2008***; ***Angilletta, 2009***). Calculating the inverse of *B*_pk_ gives an estimate of the minimum generation time (*t*_gen_) of a microbial species or strain.

As ***Smith et al.*** (***2019***) have shown, in prokaryotes, *B*_pk_ tends to increase with *T*_pk_ until ≤ 45°C, after which it levels off. To quantify this relationship in a continuous manner, we fitted a few alternative models to their dataset. More precisely, we specified ln(*B*_pk_) as the response variable, whereas the predictor was *T*_pk_, either using a natural logarithm transformation or a square root transformation. We note that we did not use a Boltzmann-Arrhenius term, even if the Metabolic Theory of Ecology framework also predicts an exponential decrease of *t*_gen_ with temperature (***Savage et al., 2004***), due to the implicit assumption that the trait being studied (i.e., *t*_gen_) is directly governed by the influence of temperature on enzyme kinetics. Such an assumption has been previously proven invalid (e.g., see ***García-Carreras et al. 2018***, ***Kontopoulos et al. 2020a***, and ***Kontopoulos et al. 2020b***). Species identity was accounted for, as a random effect on the intercept. For each model, we fitted both a non-phylogenetic and a phylogenetically-corrected variant, in the latter case using the relative chronogram of ***Kontopoulos et al. (2020b)***. All models were fitted with MCMCglmm, by running two independent chains per model for five million generations, with parameter samples being obtained from the posterior distribution every thousand generations. Samples from the first 500,000 generations were treated as burn-in and were discarded. Convergence was examined as previously described. The best-fitting model was determined according to DIC. To examine the proportion of variance explained by the fixed effects alone (Var_fixed_), by the random effect (Var_random_), or left unexplained (Var_resid_), we calculated the marginal and conditional *R*^2^ values (***Nakagawa and Schielzeth, 2013***):

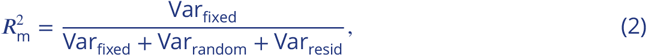

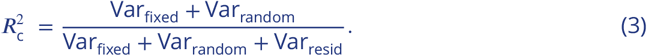

The results are shown in Appendix 1—table 5 and Appendix 1—figure 14.

**Appendix 1—table 5.**
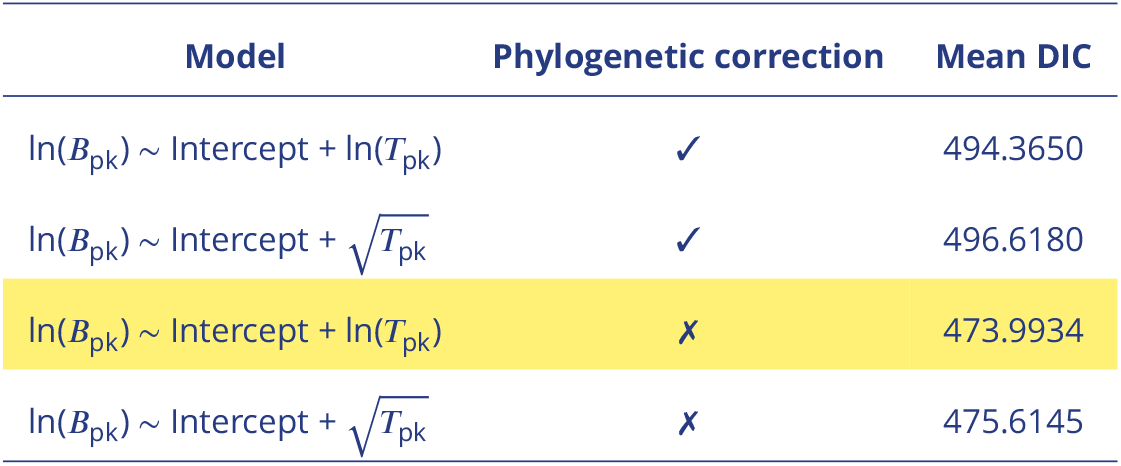
Alternative models for the relationship of fin(*B*_pk_) with *T*_pk_ across prokaryotes. The model with the lowest DIC is highlighted.

| Model | Phylogenetic correction | Mean DIC |
| --- | --- | --- |
| $\ln(B_{\text{pk}}) \sim \text{Intercept} + \ln(T_{\text{pk}})$ | ✓ | 494.3650 |
| $\ln(B_{\text{pk}}) \sim \text{Intercept} + \sqrt{T_{\text{pk}}}$ | ✓ | 496.6180 |
| $\ln(B_{\text{pk}}) \sim \text{Intercept} + \ln(T_{\text{pk}})$ | ✗ | 473.9934 |
| $\ln(B_{\text{pk}}) \sim \text{Intercept} + \sqrt{T_{\text{pk}}}$ | ✗ | 475.6145 |

**Appendix 1—figure 14.**
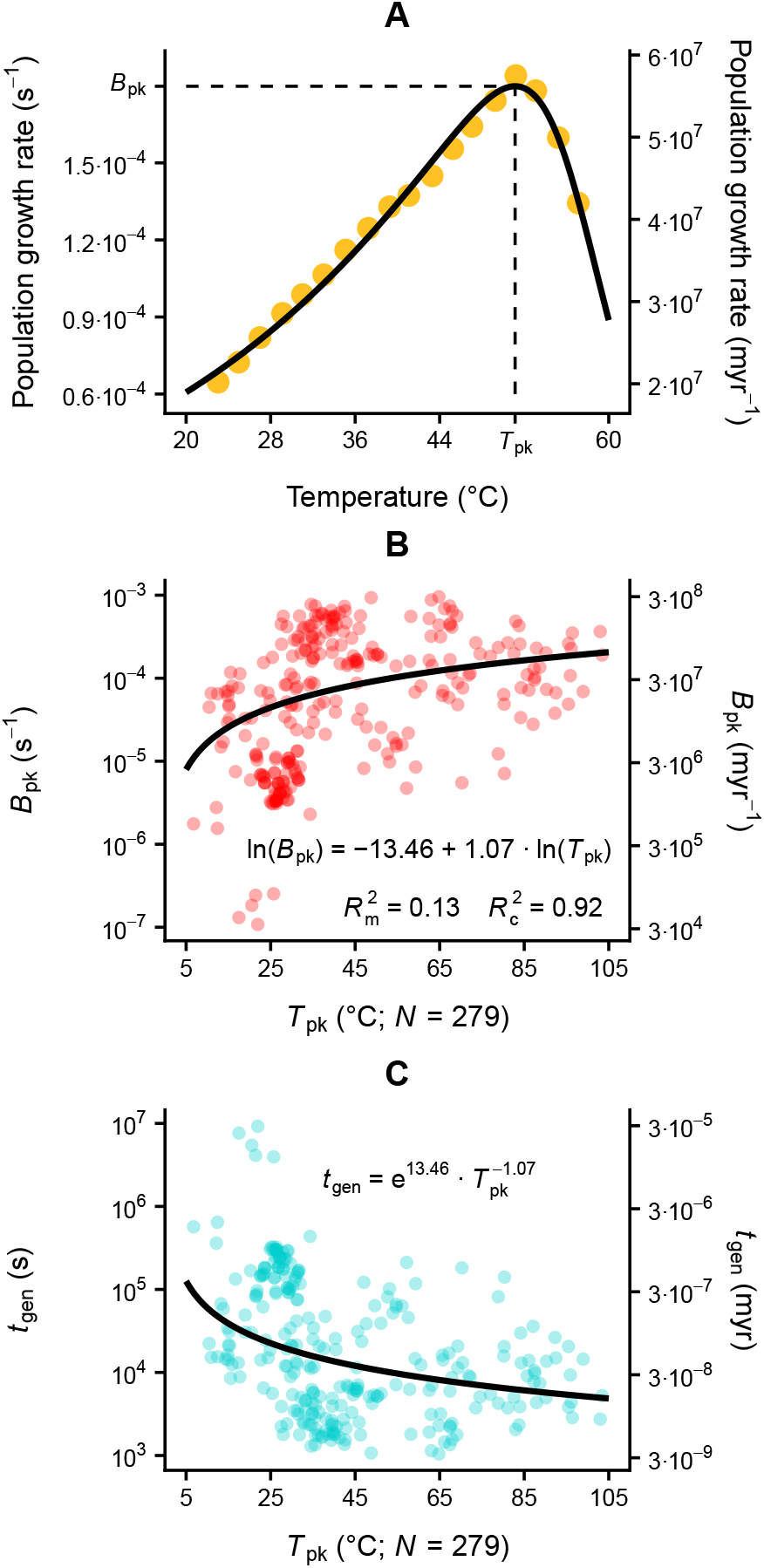
The relationships of maximum population growth rate (*B*_pk_) and minimum generation time (*t*_gen_) with *T*_pk_. A: The thermal performance curve of population growth rate of the archaeum *Natrinema pellirubrum* (***Robinson et al., 2005***). Gold circles stand for the experimental measurements, whereas the black line is the fit of the four-parameter variant of the four-parameter Sharpe-Schoolfield model (***Schoolfield et al., 1981***). *T*_pk_ is the temperature at which growth rate takes its maximum value, *B*_pk_. The left vertical axis is in units of s−1, whereas the right vertical axis is in myr−1. B: In prokaryotes, *B*_pk_ increases with the natural logarithm of *T*_pk_. This explains 13% of variance, whereas species identity leads to 92% of the variance being explained. The equation that is shown has *B*_pk_ in units of s−1 and *T*_pk_ in °C. C: *t*_gen_ decreases as a power law of *T*_pk_.

## X.II Cell volume

We further examined whether cell volume also influences the minimum generation times of prokaryotes. To this end, we estimated cell volumes for as many species in the dataset of ***Smith et al.*** (***2019***) as possible. We then fitted models with MCMCglmm that had fin(*B*_pk_) as the response and ln(*T*_pk_) and/or ln(cell volume) as fixed effects. Species identity was again specified as a random effect on the intercept, both with and without a phylogenetic correction. Given that the cell volume of most species was available as ranges and not as a single value, we incorporated the uncertainty in our models by fitting them 30 times. For each of the thirty runs, we set each species’ volume by sampling from a uniform distribution with bounds equal to the minimum and maximum cell volume of the species. Two independent MCMC chains were run for each model, for two million generations. Samples from the posterior were obtained every thousand generations after the first two hundred thousand generations and were examined for suZcient convergence. Model selection was done according to DIC, averaged across the thirty runs.

Models with phylogenetic correction had higher DIC values than their non-phylogenetic equivalents. Across the latter, the 95% HPD interval of the coefficient of ln(cell volume) almost always included zero. This was the case both when ln(cell volume) was the only fixed effect (Appendix 1—figure 15A) and when it was accompanied by ln(*T*_pk_) (Appendix 1—figure 15B). This shows that cell volume does not systematically influence *B*_pk_ across prokaryotes.

**Appendix 1—figure 15.**
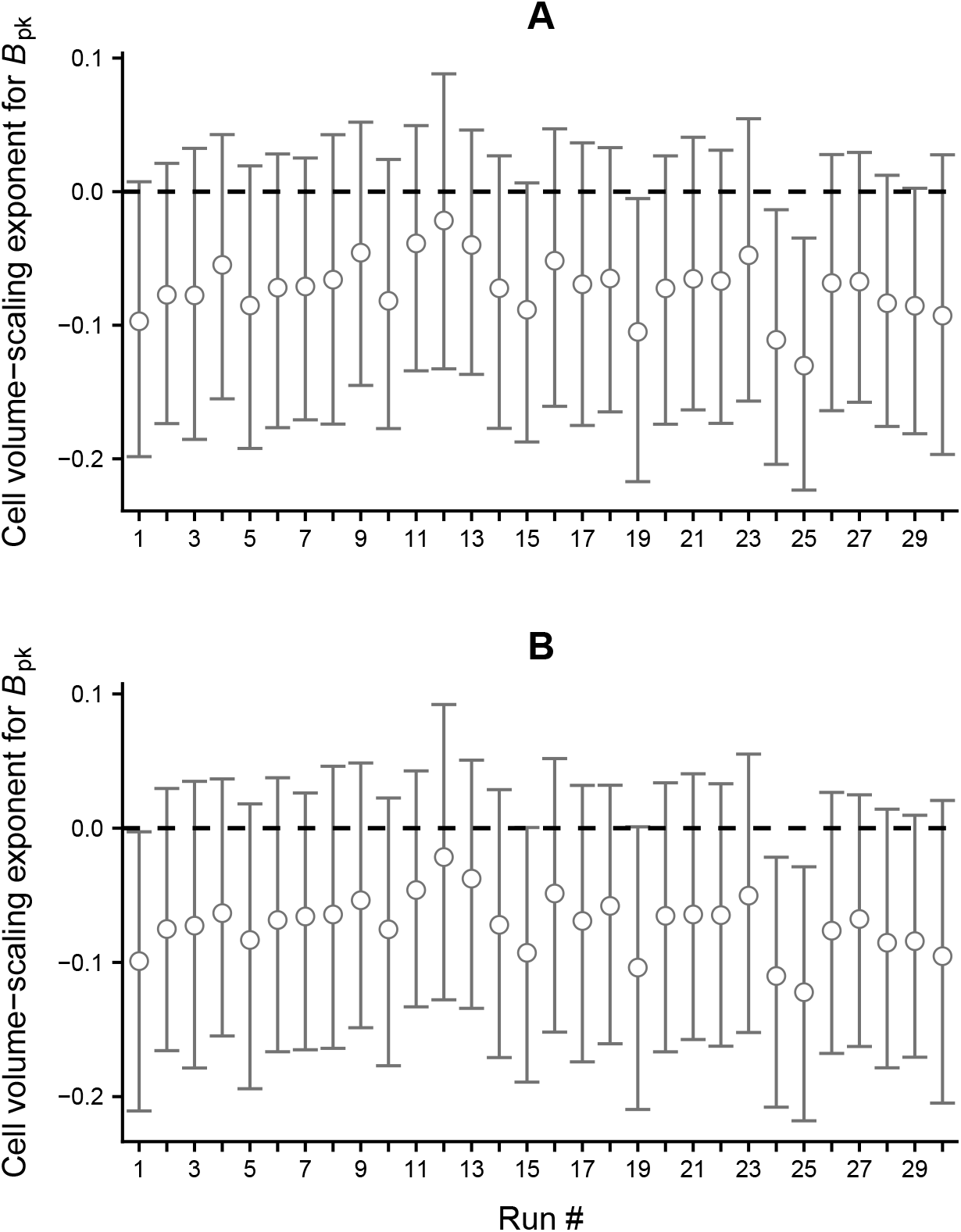
Investigation of a possible effect of cell volume on *B*_pk_ and, by extension, on generation time. Circles show the mean of the posterior distribution of the exponent of cell volume, whereas whiskers indicate the 95% HPD interval. The estimates in panel A were obtained through a regression of ln(*B*_pk_) against the natural logarithm of cell volume. In contrast, in panel B, the regression also contained the natural logarithm of *T*_pk_ as another 1xed effect.

## XI. The relationship between substitution rate and temperature

According to the neutral theory of molecular evolution (***Kimura, 1987***), the rate of molecular substitution per time (*K*) can be predicted by the generation time (*t*_gen_), the mutation rate per generation (*U*), and the proportion of selectively neutral mutations (*f*_0_):

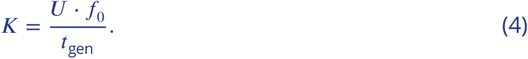

This equation makes the assumption that selectively beneficial mutations are extremely rare and can be ignored, i.e., 1 − *f*_0_ stands for the proportion of deleterious mutations. Temperature influences *t*_gen_ (Appendix 1—figure 14), but also potentially the numerator of Appendix 1—equation (4), given that mutations become more detrimental with temperature as we show in this study. Thus, *f*_0_ and possibly *U* should decrease with temperature. In contrast, ***Gillooly et al.*** (***2005***, 2007) have argued that, similarly to mass-specific metabolic rate, *U* should increase with temperature and associate (positively in prokaryotes, negatively in protists and metazoans; ***DeLong et al. 2010***) with body size (or cell volume). Such a pattern would be consistent with the “metabolic rate hypothesis” which posits that mutation rate depends on metabolic rate, due to the production of oxidative free radicals as a byproduct of metabolism. Thus, to estimate the potential effects of cell volume (*V*) and temperature (*T*) on the numerator of Appendix 1—equation (4), the equation can be written as

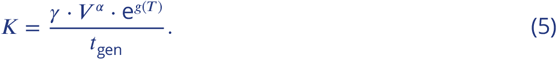

Here, *γ* is a species-specific normalisation constant that is independent of cell volume and temperature, *α* is a dimensionless exponent, whereas *g*(*T*) a temperature dependence function for which we used either a logarithmic (*g*(*T*) = *ζ* · ln(*T*)) or a quadratic equation (*g*(*T*) = *η* · *T* + *θ* · *T*^2^). It is worth noting that we did not model temperature as a Boltzmann-Arrhenius function (***Gillooly et al., 2007***) because the latter implies that substitution rates are directly governed by the effects of temperature on the activity of underlying enzymes. This assumption has been previously shown to be invalid for metazoans (***Lanfear et al., 2007***).

Appendix 1—equation (5) can be simplified by removing *T* and/or *V* if these are found to have no systematic effect on *K*. To understand if and how temperature drives substitution rates, we multiplied both sides of Appendix 1—equation (5) with *t*_gen_ and took the natural logarithm of both sides:

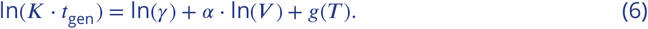

The mean DIC values of alternative fitted models are shown in Appendix 1—table 6. Models which included the natural logarithm of cell volume as a fixed effect are not shown, as the coefficient of cell volume always had a 95% HPD interval that included zero.

**Appendix 1—table 6.** Candidate models for the estimation of generation time-corrected substitution rates. The model with the lowest DIC is highlighted.

| Model | Phylogenetic correction | Mean DIC |
| --- | --- | --- |
| $\ln(K \cdot t_{\text{gen}}) \sim \text{Intercept}$ | ✓ | 161.3176 |
| $\ln(K \cdot t_{\text{gen}}) \sim \text{Intercept} + \ln(\text{Temperature})$ | ✓ | 29.3025 |
| $\ln(K \cdot t_{\text{gen}}) \sim \text{Intercept} + \text{Temperature} + \text{Temperature}^2$ | ✓ | 15.2959 |
| $\ln(K \cdot t_{\text{gen}}) \sim \text{Intercept}$ | ✗ | 188.1174 |
| $\ln(K \cdot t_{\text{gen}}) \sim \text{Intercept} + \ln(\text{Temperature})$ | ✗ | 24.9966 |
| $\ln(K \cdot t_{\text{gen}}) \sim \text{Intercept} + \text{Temperature} + \text{Temperature}^2$ | ✗ | 27.1478 |

## XII. Literature data on the effects of temperature on mutation rate

**Appendix 1—figure 16.**
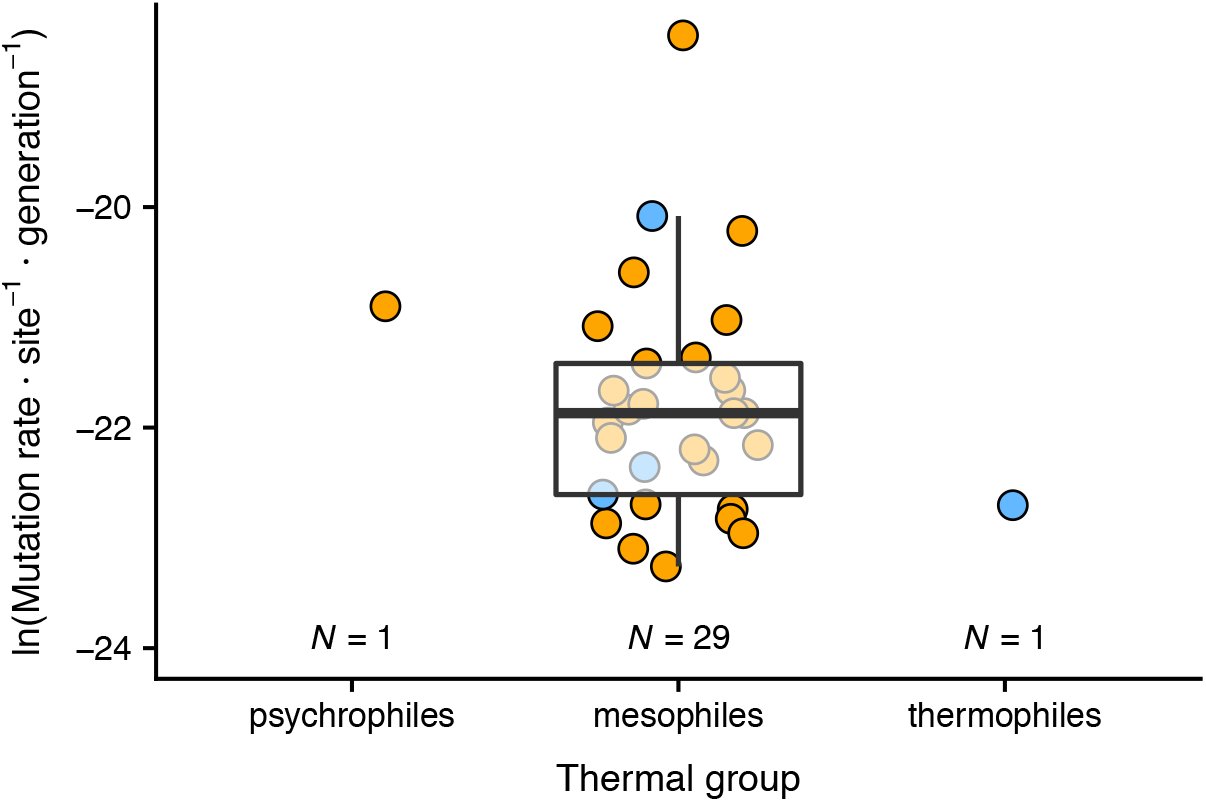
Mutation rate measurements for bacteria adapted to different thermal environments. Each data point corresponds to a different species, whereas colors represent the experimental approach. Mutation accumulation experiments (currently the most accurate method; ***Gibson et al. 2018***) are shown in orange. Boxplot edges represent the first and third quartiles, whereas the solid line stands for the median. Whiskers extend up to the most remote data point within 1.5 interquartile ranges from each boxplot edge. Most measurements are for species adapted to mesophilic environments and, therefore, no conclusion on the effect of temperature can be drawn. Data points were obtained from ***Lynch et al. (2016)***; ***Gibson et al.*** (***2018***); ***Long et al. (2018)***; ***Gibson and Eyre-Walker (2019)***.

